# STARComm Scalably Detects Emergent Modules of Spatial Cell-Cell Communication in Inflammation and Cancer

**DOI:** 10.1101/2025.08.07.669133

**Authors:** Khoa L. A. Huynh, Bruno F. Matuck, Deiziane de Souza, XiuYu Zhang, Giancarlo Fatobene, Luiz Alberto Valente Soares Junior, Luiz Fernando Ferraz da Silva, Vanderson Geraldo Rocha, Kevin M. Byrd, Jinze Liu

**Affiliations:** Department of Biostatistics, Virginia Commonwealth University, Richmond, VA, USA; Department of Oral and Craniofacial Molecular Biology, Philips Institute for Oral Health Research, Virginia Commonwealth University, Richmond, VA, USA; Department of Pathology, Medicine School of University of Sao Paulo, SP, BR, Sao Paulo, BR; Laboratório de Investigação Médica (LIM) 31, Hospital das Clínicas HCFMUSP, Faculdade de Medicina, Universidade de São Paulo, Brazil; Division of Dentistry of Hospital das Clinicas of University of Sao Paulo, SP, BR, Sao Paulo, BR; Massey Comprehensive Cancer Center, Virginia Commonwealth University, Richmond, VA, USA

**Keywords:** Spatial Transcriptomics, Cell-Cell Communication, Systems Biology, Cellular Neighborhoods, Cancer, Tumor Microenvironment, Inflammation, Precision, Medicine, Computational Pathology

## Abstract

In humans, cell-cell communication orchestrates tissue organization, immune coordination, and repair, yet spatially mapping these interactions remains a challenge for biology. We introduce STARComm, a scalable-interpretable computational method that identifies Multicellular Communication Interaction Modules (MCIMs) by detecting spatially co-located receptor-ligand activity from high-plex spatial transcriptomics in 2D and 3D. Applied to an atlas of >14million cells across 8 cancers, STARComm revealed 24 conserved and tumor-specific MCIMs, including a fibro-immune module with targetable axes linked to immune exclusion and immunotherapy resistance. In chronic graft-versus-host disease, STARComm identified three salivary gland MCIMs predictive of patient death and two druggable axes (*CXCL12-CXCR4*, *CCL5-SDC4*), both with FDA-approved therapeutics. STARComm demonstrated that peripheral tissue profiling can forecast fatality nearly 3 years in advance using minor salivary glands. By enabling scalable biomarker discovery, drug targeting, and spatially resolved precision profiling, STARComm bridges the gap between spatial biology and clinical translation, advancing the field of spatial medicine.

**SUMMARY:** Despite major advances in spatial biology, no framework has yet linked spatially resolved intercellular communication networks, independent of cell types, to clinical outcomes in human disease. Here, we present STARComm, a scalable method that identifies Multicellular Interaction MCIMs (MCIMs). Applying STARComm to minor salivary gland biopsies from patients with chronic graft-versus-host disease (GVHD), we identify MCIMs that not only distinguish healthy from diseased tissue but also stratify patient survival. High-risk MCIMs are enriched for actionable immune and stromal pathways, including those targetable with existing therapies. These findings establish the first outcome-linked spatial communication framework in any human disease and highlight the translational potential of oral tissues as minimally invasive platforms for real-time immune diagnostics, prognostic modeling, and therapeutic screening.

## INTRODUCTION

Chronic graft-versus-host disease (GVHD) remains the most common cause of late non-relapse mortality following allogeneic hematopoietic cell transplantation, affecting 30%–70% of transplant recipients who survive beyond 100 days^1,2^. Since the 1950s, nearly 2 million HCTs have been performed^3^; among these, up to 40%, including nearly 1 million individuals globally, may have experienced moderate to severe GVHD^4,5^. Despite decades of clinical observation, no spatially resolved or serum biomarkers are clinically approved to predict GVHD-related fatality, and current diagnostic approaches are often imprecise, lack sensitivity at early stages or may require tissue not easily accessible for longitudinal monitoring^6–13^. Given the disease’s rarity, clinical heterogeneity, and systemic manifestations, there is a pressing need for tools that can detect early signs of immune dysregulation in accessible peripheral tissues and biofluids, ideally in a format scalable across patients, organs, and disease contexts.

At the core of GVHD pathogenesis is dysregulated immune communication^14^. The adult body maintains tissue homeostasis through quadrillions of cell–cell interactions, primarily mediated by receptor–ligand (R–L) signaling. Single-cell RNA sequencing (scRNA-seq) has enabled the inference of such communication events via tools like CellChat, NicheNet, CellPhoneDB, LIANA+, NATMI, and scSeqComm to infer cell-cell communication; however, these methods share two major limitations^15–20^. First, they typically infer communication between broad, predefined cell types, overlooking the diversity of interactions at single-cell resolution. Second, they lack spatial context—making it impossible to confirm whether potentially interacting cells are physically proximal within the tissue architecture. These limitations lead to high false-positive rates and limit our ability to capture emergent communication patterns within intact tissues^7–9^. This and current clinical limitations of scRNAseq make using available atlases of GVHD less relevant in the era of digital biology and spatial medicine^21–23^.

Spatial transcriptomics technologies provide a deeper understanding of cell-cell communication (C2C) by mapping gene expression within their native tissue microenvironments^24^. A variety of tools have been developed for spatial communication analysis, including SpatialDM, SpaTalk, COMMOT, and CellNEST, each offering unique innovations, from bivariate spatial autocorrelation to knowledge graphs and optimal transport^25–28^. However, a fundamental challenge arises because most of these tools were designed for two-dimensional and spot-level spatial transcriptomics^26^. Overall, each is limited by either their resolution, scalability, and/or capacity to capture single-cell-level signaling heterogeneity. As high-resolution spatial multiomics technologies mature, new methods are needed to handle their scale, sparsity, and complexity.

To address these challenges and potential application opportunity for spatial medicine in GVHD, we developed STARComm (<u>S</u>pa<u>TiA</u>l <u>R</u>eceptor–Ligand Analysis for Cell–Cell <u>COMM</u>unication), a scalable and unsupervised algorithm that identifies a new concept a la tissue cellular neighborhoods^29,30^, called Multi-Cellular Interaction MCIMs (MCIMs), which are emergent spatial neighborhoods defined by collective R–L signaling. STARComm operates in two phases: global detection of all active R–L interactions across tissues, and spatial clustering of co-localized signals into MCIMs. This global-to-local strategy enables discovery of complex interaction patterns across millions of cells, in 2D and 3D datasets, without requiring prior cell-type annotation. We benchmarked STARComm using simulated spatial datasets and healthy tissue atlases spanning six oral and glandular tissues (>1.2 million cells), where it outperformed existing methods in speed, scalability, and accuracy. Applying STARComm to eight solid tumor types uncovered conserved and disease-specific MCIMs linked to immune evasion, neoangiogenesis, and fibroblast–immune suppression—many of which are already targeted by approved or investigational drugs, highlighting its translational potential for immune-oncology therapies. In autoimmune graft-versus-host disease, STARComm identified specific signaling modules in salivary glands that predicted long-term survival, with certain modules involving targetable axes like CXCL12–CXCR4 and CCL5–SDC4, providing prognostic biomarkers nearly three years in advance. Using spatial-Drug2Cell, we mapped FDA-approved drug targets within these fatality-linked MCIMs, demonstrating the potential of this approach for localized therapeutic inference^31^.

This work highlights how minimally invasive sampling of peripheral tissues such as salivary glands can uncover spatial immune circuits that stratify systemic risk—the first example of this for spatial biology that has been published and broadening the definition of oral-systemic linkage long reported^32,33^. Furthermore, in this study, MCIMs serve not only as biomarkers of prognosis but also as guides for therapeutic interception, drug design, response prediction, and toxicity monitoring. By enabling the direct linkage of spatial cell–cell communication patterns to clinical outcomes, STARComm and the concept of MCIMs introduce a generalizable biomarker framework for immune dysregulation with broad applicability across GVHD, autoimmunity, infection, and cancer.

## RESULTS

### Mapping Clinically Relevant Tissue Communication MCIMs with STARComm

Chronic GVHD, like many immune-mediated diseases, stems from systemic yet tissue-specific immune dysregulation; however, the spatial organization of pathogenic signaling events remains poorly understood^34^. Central to this pathology is cell–cell communication (C2C), a process rarely driven by a single receptor–ligand (R–L) pair in time or space. Instead, multiple signaling pathways operate concurrently within a shared tissue microenvironment, forming complex interaction landscapes (Figure 1a). To bring biological and spatial structure to this known complexity, we formulated the concept of Multi-Cellular Interaction (MCIMs), which are like tissue cellular neighborhoods or described tissue “ecotypes”^29,35^. These C2C hotspots of spatially coordinated signaling represent emergent functional units of multicellular cooperation (Figures 1b-d). MCIMs capture the integrated behavior of diverse cell types and R–L pairs as they orchestrate homeostasis, immunity, and disease progression. We hypothesized that mapping these spatially organized signaling MCIMs would reveal conserved architectures underlying chronic inflammation and help stratify clinical outcomes in GVHD and beyond.

**Figure 1.**
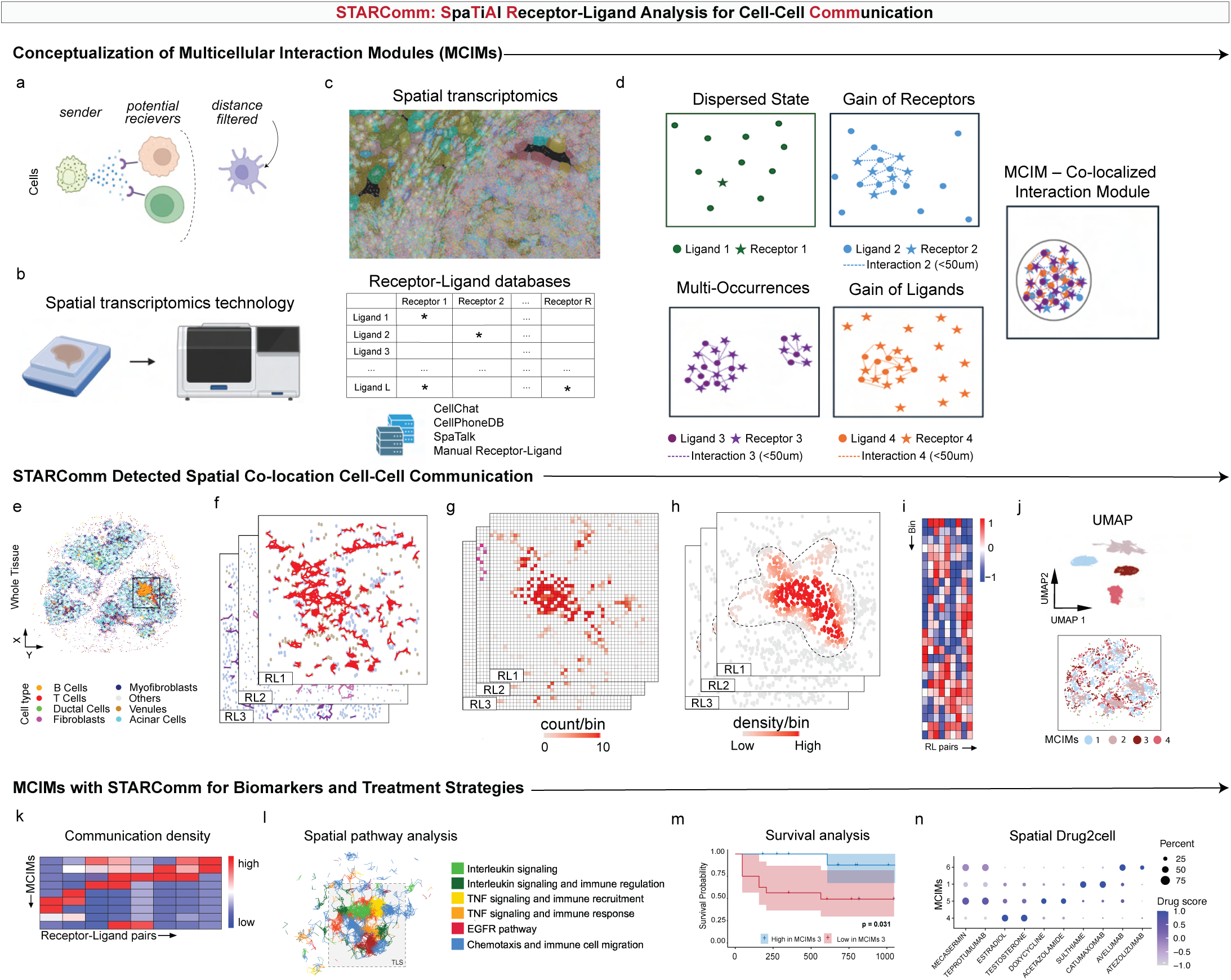
Workflow of SpaTiAl Receptor-Ligand Analysis for Cell-Cell Communication (STARComm). | **a**, Conceptual framework: cell–cell communication is inferred based on spatial proximity, where ligand-expressing (sender) cells are more likely to interact with nearby receptor-expressing (receiver) cells. **b**, Tissue samples are collected and processed using spatial transcriptomics (e.g., Xenium). **c**, Spatial gene expression images are generated, with cell type annotations (top) and receptor–ligand pairs curated from public databases (e.g., CellChat, CellPhoneDB, SpaTalk) or custom lists (bottom). **d**, Communication is modeled across various scenarios: non-interacting dispersed states, gain of receptor expression, gain of ligand expression, or recurring local interactions. A multicellular communication interaction module (MCIM) is defined as a set of co-localized receptor–ligand interactions. **e**, Spatial transcriptomics data with annotated cell types visualized by color. **f**, Communication edges are inferred by identifying ligand-expressing and receptor-expressing cells using TACIT and then connecting cells within a 50µm spatial radius. **g**, The tissue is partitioned into spatial bins; receptor–ligand edges are counted per bin. **h**, Kernel density estimation generates a spatial map of communication density for each receptor–ligand pair. **i**, Communication densities are summarized in a matrix (bins × receptor–ligand pairs), where values represent interaction densities. **j**, This matrix is clustered to identify MCIMs. **k**, Heatmap of communication densities for each MCIM. **l**, MCIMs are grouped by signaling pathway and mapped onto spatial tissue architecture. **m–n**, MCIMs can be associated with clinical outcomes, such as survival, and may serve as biomarkers or therapeutic targets (e.g., in GVHD).

To identify and characterize these MCIMs in spatial data, we developed STARComm, a computational framework with a robust three-step process. First, the STARComm workflow begins by identifying high-confidence communication edges between individual cells. This step defines potential interactions by locating signal-sending (ligand-expressing) and signal-receiving (receptor-expressing) cells based on both their gene expression levels and spatial proximity. To ensure robustness, we adopt the optimal thresholding algorithm from TACIT, which establishes specific, context-aware expression cutoffs for each ligand and receptor gene (Figure 1e-f)^36^. Cells exceeding these dynamic thresholds are designated as positive, minimizing false positives from background noise or data sparsity. A communication edge is then established between a positive ligand-cell and a corresponding positive receptor-cell if they are located within a defined spatial distance, generating a comprehensive map of connections for each individual receptor-ligand pair. In the second step, these discrete communication edges are transformed into continuous spatial density maps. The tissue space is first discretized using a grid, and the local abundance of each type of interaction is quantified using kernel density estimation (KDE). To accurately position each interaction, we use the midpoint between the centers of the sending and receiving cells (Figure 1g). The KDE process converts the discrete interaction points into a smooth, continuous density map for each ligand-receptor pair, where elevated values indicate regions with a high concentration of that specific signaling event (Figure 1h).

Finally, the third step integrates these individual density maps to discover the MCIMs. To enable robust comparison across different interactions and tissues, the density maps are normalized via a z-norm (Figure 1i). These normalized maps are then concatenated into a single feature matrix where rows represent spatial grid bins and columns represent all L-R pairs, thus capturing the complete communication profile for every location in the tissue. An unsupervised clustering algorithm is then applied to this matrix, grouping together spatial bins with similar communication profiles to reveal the MCIMs (Figure 1j-k). By engineering STARComm to detect and cluster spatially co-localized receptor–ligand interactions, we establish a scalable framework for resolving multicellular signaling MCIMs at single-cell resolution. This positions STARComm to uncover tissue-scale immunologic architectures that cannot be inferred from cell-type annotations or single-pair interactions alone. In the context of immune-mediated diseases like GVHD, STARComm lays the foundation for linking spatial communication structure to emergent immune dysfunction, clinical outcomes, and therapeutic vulnerabilities (Figure 1l-m).

### Benchmarking STARComm for Detecting Emergent Signaling MCIMs in Spatial Tissue Context

Since our goal was moving a biomarker like this to clinic, robust detection metrics of spatial cell–cell communication remain essential. However, the absence of ground truth datasets in biological tissues has limited the ability to systematically validate spatial communication frameworks. To overcome this barrier, we developed a simulation platform that emulates the spatial architecture, cellular density, and gene expression characteristics of high-resolution spatial transcriptomic data (e.g., MERSCOPE, Xenium), enabling direct performance comparisons across computational methods (Figure. 2a)^18^. Each simulated MCIM was defined by a specific set of receptor-ligand pairs, communication density, and expression levels, with gene expression modeled using a mixture of negative binomial distributions to represent communicating and non-communicating (Figure 1b–See Methods).

**Figure 2.**
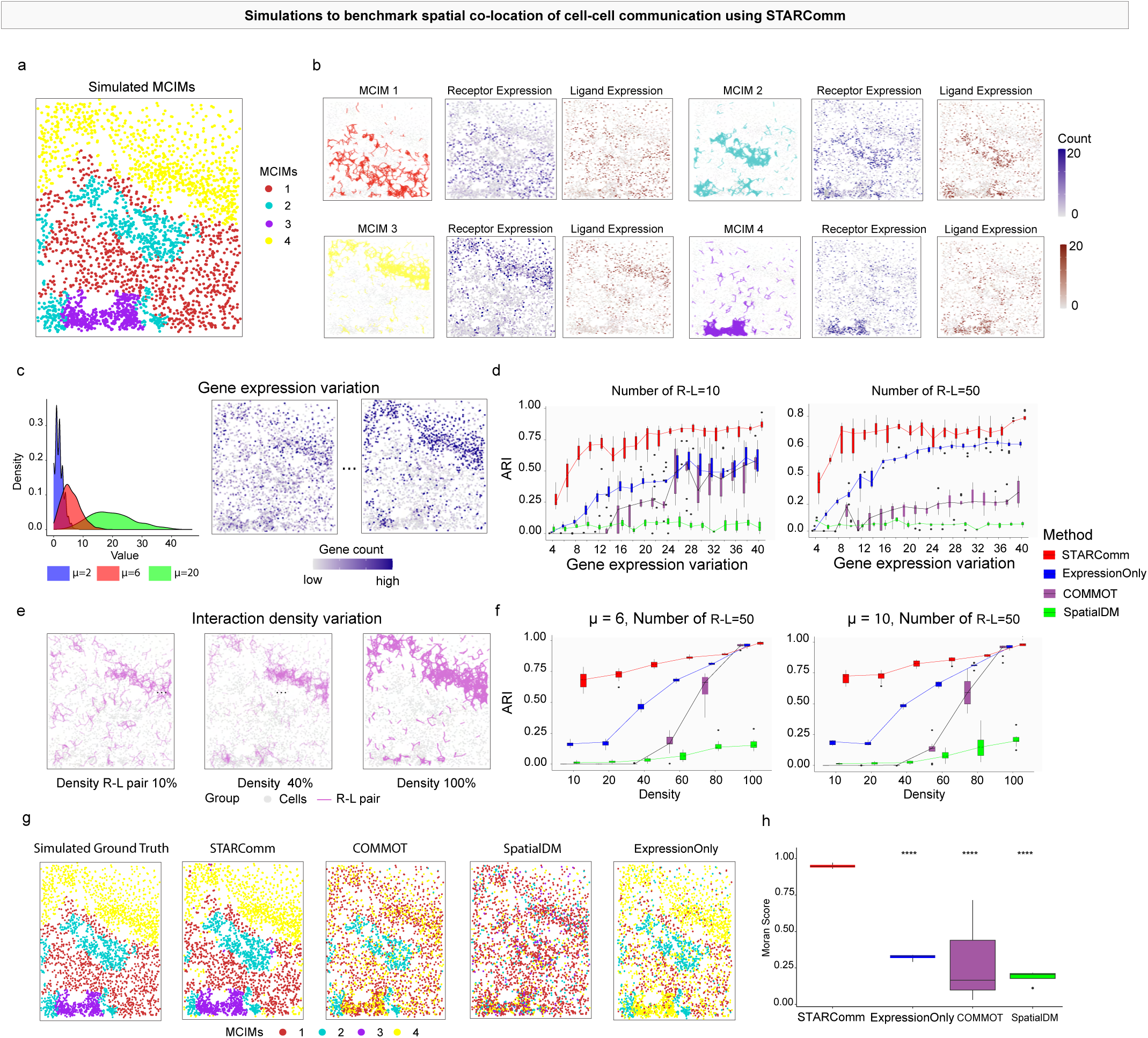
Benchmarking of STARComm Using Simulated Data. | **a**, Simulated datasets were generated by modeling spatially localized cell–cell communication events. Four distinct MCIMs were simulated, each defined by a unique set of ligand–receptor interactions serving as ground truth. **b**, A representative example of the simulated tissue illustrates ground-truth MCIMs, showing receptor–ligand expression and corresponding communication edges. **c**, For each ligand– receptor pair within an MCIM, expression values were drawn from negative binomial distributions, with communicating cells assigned high mean expression (𝜇_𝐻_) and non-communicating cells assigned low expression (𝜇_𝐿_). **d**, Benchmarking across a range of expression levels (𝜇_𝐻_ from 4 to 40) shows that STARComm achieves consistently higher adjusted Rand index (ARI) scores compared to ExpressionOnly, COMMOT, and SpatialDM, particularly under low-expression conditions. **e**, Communication density is defined as the proportion of positive communication edges among all cell pairs within a 50 μm radius. **f**, When varying communication densities from 10% to 100% across 50 high-expression ligand–receptor pairs, STARComm maintained superior ARI performance and was uniquely capable of recovering MCIMs even at the lowest density of 10%. **g**, Spatial visualizations comparing predicted communication regions with ground truth further highlight STARComm’s superior spatial localization accuracy. **h**, Moran’s I scores demonstrate significantly stronger spatial co-localization of ligand–receptor pairs using STARComm, underscoring its effectiveness in capturing spatial communication neighborhoods.

We first assessed STARComm’s ability to identify MCIMs across varying gene expression signal-to-noise ratios and complexities, benchmarking it against COMMOT, SpatialDM, and a baseline algorithm (ExpressionOnly) that performs clustering of based on only the expression level of ligand receptors without spatial information. In this experiment, the mean expression for non-communicating cells was fixed while the expression level for communicating cells was varied (Figure 1c). Across all scenarios, which included using 10, 20, 50, or 100 receptor-ligand pairs, STARComm consistently achieved the highest Adjusted Rand Index (ARI) score. Notably, at the lowest signal-to-noise ratio where other methods failed, only STARComm (mean-ARI=0.40, SD=0.13) effectively captured the ground truth MCIMs (Figure. 2d, Extended Data 1a). This ability to resolve low-abundance, spatially organized signaling is essential for capturing early immune dysfunction in chronic GVHD, where pathogenic interactions may be both rare and regionally restricted.

We then investigated performance as a function of communication density, varying the probability of communication from 10% to 100% in each MCIM (Figure 1e). STARComm again demonstrated superior performance, maintaining a high ARI of 0.7 at the lowest communication density (10%). This significantly outperformed ExpressionOnly (ARI=0.20), COMMOT (ARI=0.01), and SpatialDM (ARI=0.01) at the same challenging density level, highlighting STARComm’s sensitivity in detecting sparse communication patterns (Figure 2f). Detecting such sparse communication is particularly critical in diseases like GVHD, where early pathogenic MCIMs may emerge subtly within otherwise normal-appearing tissue and guide both risk stratification and timely therapeutic intervention.

In a third test, we next validated the spatial coherence, computational performance, and parameter stability of the results. Visual inspection of the spatial plots confirms that STARComm better captures the co-localization of R-L pairs compared to other methods (Figure 2g). We quantified this observation using Moran’s I, a measure of spatial autocorrelation, which showed STARComm’s MCIMs were significantly more spatially coherent (mean-Moran’s I=0.94) than those from ExpressionOnly, SpatialDM, and COMMOT (Figure 2h). In terms of computational efficiency, STARComm (mean-12.63sec, SD=10sec) had the shortest runtimes (Extended Data 1b). Finally, we confirmed that STARComm’s performance is highly stable across a wide range of its key parameter settings, including the number of R-L pairs, different bandwidths and grid sizes (Extended Data 1d). This combination of accuracy, speed, and robustness is vital for clinical use, enabling reliable, scalable diagnostics and real-time treatment decisions in GVHD and related disorders.

### MCIM Atlas Across Tumor Types Identifies Conserved and Targetable Communication Networks

STARComm test on a large MERSCOPE atlas of 16-samples (n=14.5 million-cells, n=500-genes, n=250 receptor-ligand pairs) across eight cancer types^19^. This identified 24-MCIMs, highlighting both unique and shared communication patterns (Figure 3a). Different cancer types show distinct communication patterns. Breast cancer mainly involves cytokine and co-stimulatory signaling, while colon cancer uniquely engages chemokine and angiogenic pathways. Liver cancer features integrin and growth factor signaling; lung cancer uses selectin adhesion and Wnt pathways; melanoma exhibits CXCL16-mediated immune recruitment. Ovarian cancer signals through T-cell co-stimulation, uterine cancer via immune escape pathways, and prostate cancer through immune regulation and cell adhesion signaling.

**Figure 3:**
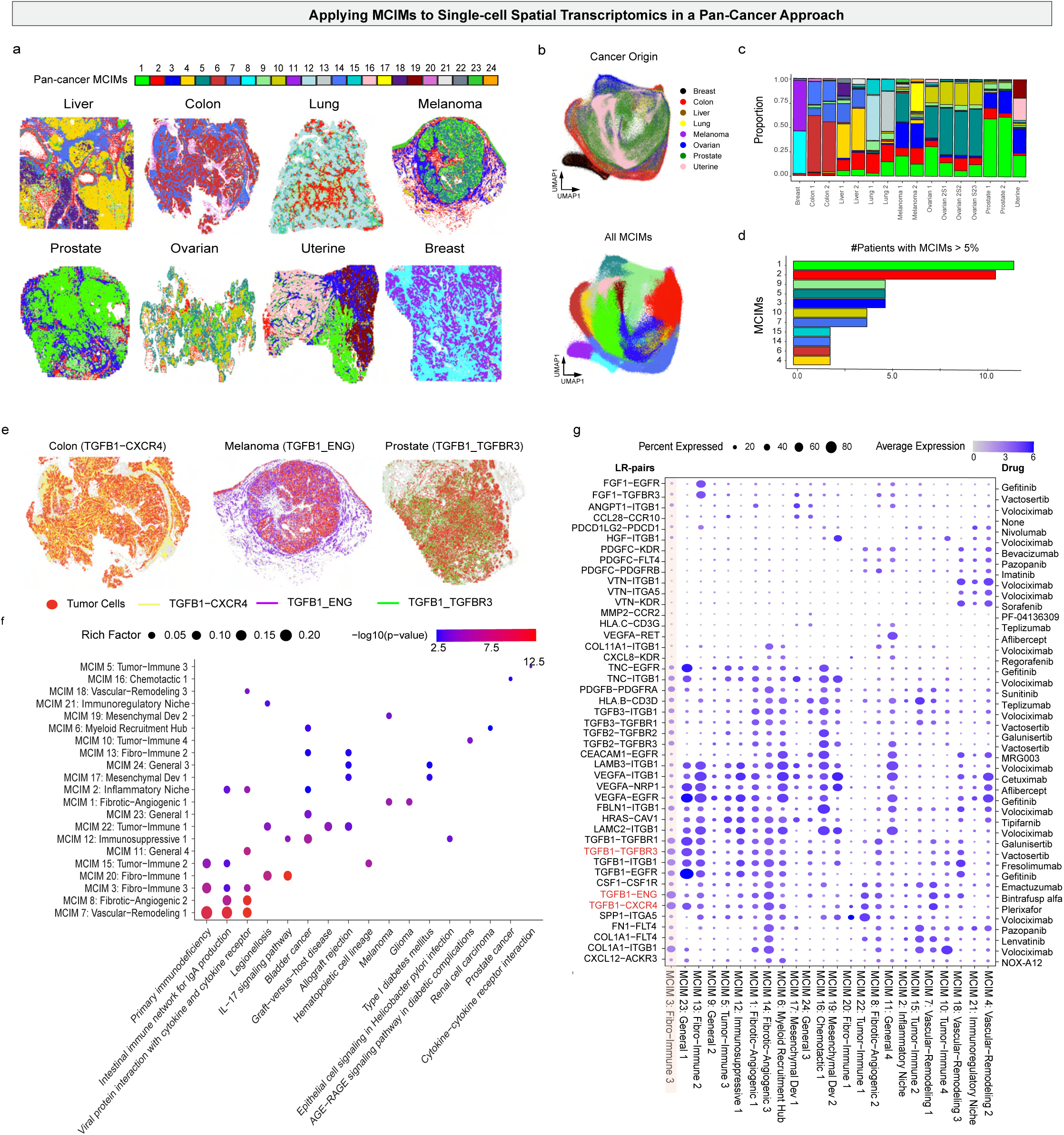
Illustrates the application of MCIM analysis to spatial transcriptomics data across eight cancer types |. **a**, The spatial maps display each cancer type with regions colored according to their associated MCIMs, highlighting distinct communication modules within the tissue. **b**, The UMAP visualization shows the density of spatial regions, with the top plot coloring regions by cancer origin and the bottom plot by MCIMs, revealing clear clustering patterns that differentiate tissue types and interaction networks. **c**, The bar plot quantifies the proportions of each MCIM across 16 tissue regions and eight cancer types, demonstrating variability in cell–cell communication patterns. **d**, The accompanying bar chart indicates the number of patients in which each MCIM accounts for more than 5% of tissue regions, emphasizing the prevalence of specific modules. **e**, Spatial maps of tumor regions highlight key receptor–ligand (R–L) interactions, such as TGFβ1–CXCR4 in colon, TGFB1-ENG in melanoma, and TGFB1-TGFBR3 in prostate cancer, illustrating localized communication pathways. **f**, KEGG pathway analysis across 24 MCIMs identifies the top three active signaling pathways, indicating biological processes driving tumor microenvironment interactions. **g**, The dot-heatmap presents MCIMs (columns) and the top three R–L pairs linked to pharmaceutical targets, revealing potential avenues for targeted therapy based on cell communication networks.

A principal component analysis (PCA) of the MCIM proportions demonstrated distinct clustering patterns associated with each cancer type (Figure 3b). For example, breast tissue exhibited high proportions of MCIMs-8 and 11, colon tissues were enriched in MCIM-6, liver in MCIM-4, lung in MCIM-15, melanoma in MCIM-3, ovarian in MCIMs-5 and 10, prostate in MCIM-3, and uterine in MCIMs-16 and 19 (Figure 3c). MCIMs-1, and 2 were the most prevalent across tissues, with MCIM-1 showing high-density communication pathways such as *CD14::ITGB1* and *TGFB1::TGFBR2*—pathways involved in immune modulation, cell proliferation, and tumor progression (Figure 3d)^25,26^. Receptor-ligand interactions within MCIMs offer promising therapeutic targets. For example, intertumoral communication pathways such as *TGFB1::CXCR4* in colon cancer, *TGFB1::ENG* in melanoma, and *TGFB1::TGFBR3* in prostate cancer are enriched and can be targeted to disrupt tumor-promoting signals (Figure 3a). MCIM4, involved in angiogenesis and ECM remodeling, could be inhibited with drugs like Bevacizumab (anti-VEGF) or Pazopanib to suppress tumor neovascularization. MCIM2, which facilitates myeloid cell recruitment through *CSF1::CSF1R* signaling and chemokine scavenging via ACKR1, might be modulated with CSF1R inhibitors such as Pexidartinib to reduce immunosuppressive myeloid infiltration^27^.

Notably, MCIM-3 emerges as a recurrent and immunosuppressive communication module across multiple cancer types—including melanoma, prostate, and uterine tumors (Figure 3, Supplement Figure 3). This module is characterized by a convergence of TGF-β signaling, CXCL chemotaxis, and leukocyte adhesion, suggesting a coordinated mechanism that excludes cytotoxic lymphocytes and promotes immune evasion^28^. Therapeutically, MCIM-3 represents a rational entry point for reprogramming “cold” tumors into “hot”. Inhibiting TGF-β (e.g., Fresolimumab) may relieve stromal-mediated suppression, while blockade of the *CXCR4::CXCL12* axis (e.g., Plerixafor) could dismantle chemokine traps and facilitate T-cell and NK-cell infiltration (Figure 3f-g)^29^. Additionally, disrupting *TGFB1::ENG* signaling, a key fibroblast– endothelial interaction enriched in MCIM-3, may attenuate the fibro-immunosuppressive niche and enhance vascular accessibility. These converging strategies highlight MCIM-3 as a dynamic functional unit—not merely a spatial co-localization, but a coordinated, targetable immune regulatory circuit. Targeting MCIM-3 may synergize with immune checkpoint inhibitors (e.g., anti– PD-1) and unlock combinatorial treatment regimens across multiple tumor contexts. By mapping emergent communication modules across diverse solid tumors, STARComm uncovers a spatial logic to the tumor microenvironment, identifying tissue-specific and pan-cancer MCIMs that serve as both mechanistic insights and actionable entry points for precision immunotherapy.

### Tissue-Scale Organization of Cell–Cell Signaling in Uterine Cancer

STARComm was validated on real uterine cancer datasets from MERSCOPE (250 receptor-ligand pairs, 843,285-cells)^19^. The tumor microenvironment displays substantial heterogeneity within individual tumors. Using STARComm on uterine cancer tissue, we identified three distinct large spatial regions, each characterized by MCIMs-1, 3, and 16 (Supplement Figure 3a-b). These MCIMs suggested that even within a single tumor, different regions were actively engaged in distinct, and potentially conflicting biological processes. MCIM-16, involving tumor, fibroblasts, B-cells, and vascular-cells, shows high *CEACAM1−EGFR* and *ICAM1−MUC1* signaling, promoting tumor growth, immune regulation, and angiogenesis^20,21^. MCIM-1, also with tumor, stromal, and immune cells, features *WNT5A−FZD7* signaling, linked to cell proliferation and migration^22^. MCIM-3, enriched in tumor and T-cells, has elevated *HGF−ITGB1* signaling, tied to cell activation, adhesion, and leukocyte recruitment, suggesting a role in inflammation and tumor development (Supplement Figure 3c-f)^23^. GO enrichment analysis shows different biological processes in MCIMs region. MCIM-1 relates to development (neuron differentiation, digestive system development), influencing tissue formation and metabolism. MCIM-3 is linked to immune functions (chemotaxis, cell activation, adhesion, leukocyte activation), indicating immune cell recruitment and inflammation in tumor progression. MCIM-16 mainly involves immune cell chemotaxis and migration, emphasizing immune cell trafficking in the tumor region (Supplement Figure 3g). Hence, STARComm identifies distinct communication zones within tumors.

### STARComm Framework for Mapping Spatial Communication at Scale: Co-location Communication in VisiumHD Colon Cancer and 3D Merscope Mouse Brain Datasets

Integrating four VisiumHD datasets (n=2,518,890 spots, n=3000 R-L pairs), we applied STARComm and observed significantly enhanced multicellular communication in cancerous regions compared to normal tissue (p=0.017)^24^(Figure 4a-c, e). Moreover, we identified MCIM 6, a distinct multicellular interaction module positioned at the tumor periphery. This specific location suggests a critical role in shaping the tumor microenvironment and influencing tumor progression (Figure 4f). MCIM-6 was characterized by communication among cancer-associated fibroblasts (CAFs), endothelial cells, tumor cells, pericytes, and macrophages, indicating a complex interplay of cell types known to modulate tumor growth, angiogenesis, and immune responses (Figure 4g). Further investigation revealed key R-L interactions within MCIM-6, including *FBN1-ITGAV*, *THBS2-ITGA6*, *MMP2-FGFR1*, *CCL14-ACKR1*, and *COL1A1/A2-FLT4*, which were implicated in matrix remodeling, angiogenesis, immune modulation, and lymphatic vessel formation (Figure 4d). The emergence of MCIM-6 exclusively at the tumor-stromal interface reveals how cancer co-opts spatially organized communication hubs to coordinate matrix remodeling, vascular invasion, and immune suppression, highlighting MCIMs as actionable units of pathological tissue reprogramming.

**Figure 4:**
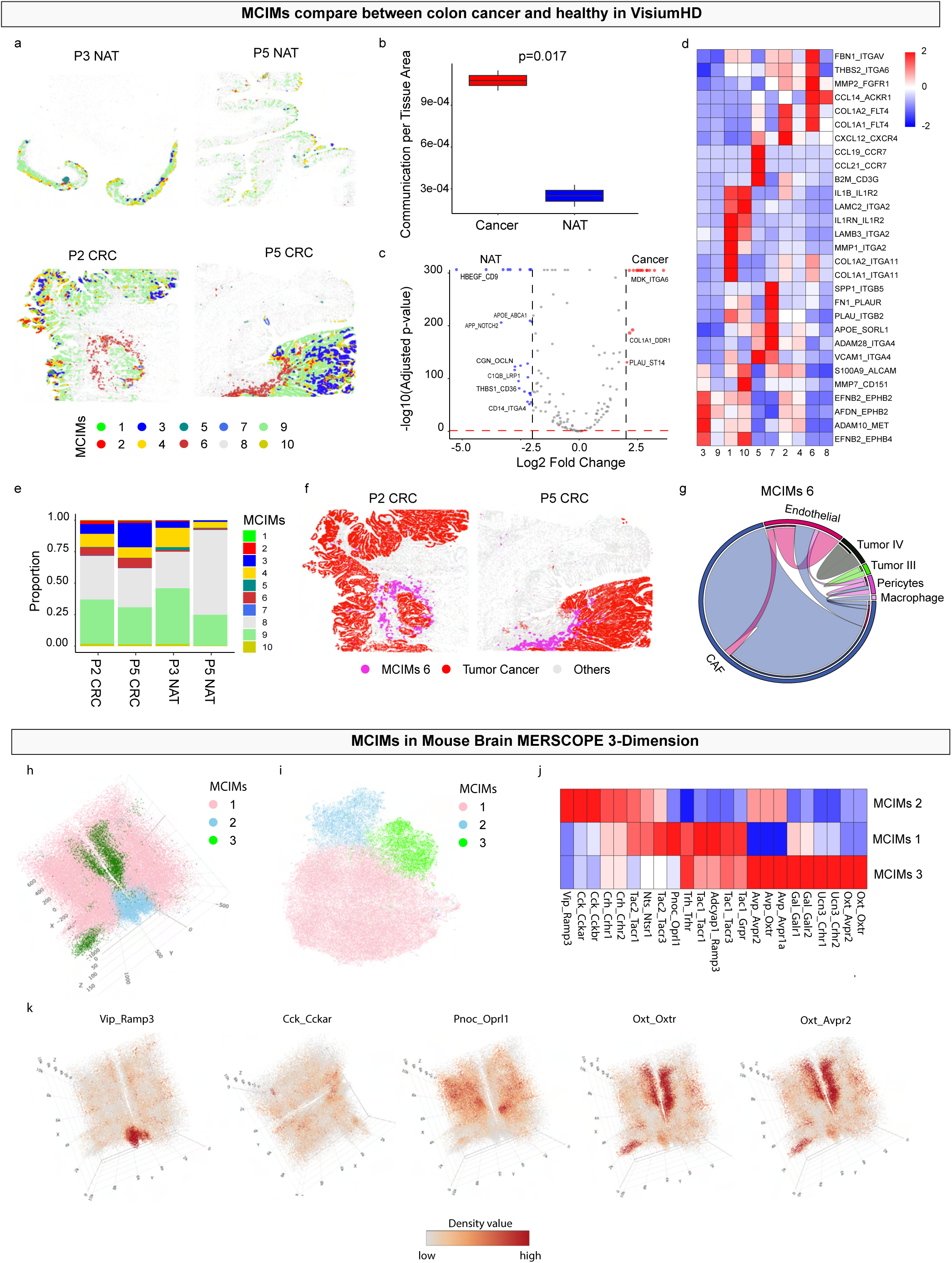
Compares multicellular interaction modules (MCIMs) between colon cancer and healthy regions within VisiumHD datasets |. **a**, Spatial maps display four tissue regions, with MCIMs color-coded to distinguish two colon cancer areas (P2 CRC and P5 CRC) and two healthy regions (P3 NAT and P5 NAT). **b**, A boxplot demonstrates a significant increase in cell–cell communication activity within the cancer tissues compared to healthy tissues (p=0.017, Wilcoxon test), indicating heightened intercellular signaling in tumors. **c**, The volcano plot highlights differentially enriched receptor–ligand pairs between healthy and cancer regions, emphasizing specific signaling interactions associated with malignancy. **d**, The heatmap portrays signature R– L pairs characterizing each MCIM, revealing distinct communication profiles across regions. **e**, Additionally, the proportion of R–L pairs varies, **f**, with MCIM 6 (pink) notably enriched in tumor areas (red), suggesting its prominent role in tumor microenvironment interactions.

We applied STARComm to a 3D MERSCOPE dataset of intact mouse brain tissue (200μm, n=133,848 cells, n=23 receptor-ligand pairs) to map spatial communication networks. This analysis revealed three distinct microdomains of coordinated intercellular messaging (MCIMs; Figure 4h-i). MCIM-1 was characterized by co-localization of Pnoc::Oprl1 and Tac2::Tacr3, MCIM-2 by Vip::Ramp3, Cck::Cckar, and Cck::Cckbr, and MCIM-3 by Oxt::Oxtr, Oxt::Avpr2, Ucn3::Crhr1, and Ucn3::Crhr2 (Figure 4j-k). This demonstrates STARComm’s versatility in analyzing spatial communication networks in both 2D and 3D tissue architectures.

### STARComm of Glandular Tissues Reveals Communication in Healthy

In our previous work, we showed that periductal inflammation in glands consisted of aggregates of B and T cells surrounded by dendritic cells and small vessels, forming an architecture consistent with TLS^36^. Across many chronic inflammatory diseases like GVHD, the presence of tertiary lymphoid structures (TLS) supports local activation and maintenance of alloreactive progenitor T cells, and the extent of TLS formation is increasingly correlated with disease severity and progression^37^. To understand how immune cells are sustained within TLS in the periductal area and throughout the parenchyma, we started by analyzing MCIMs in healthy tissues. We applied STARComm to high-resolution spatial transcriptomics data generated through the Human Cell Atlas Oral and Craniofacial Bionetwork^30,38^. We used sequential sections from well-annotated spatial blocks already used for RNAscope, MERSCOPE (sp-transcriptomics) and Phenocycler-Fusion 2.0 (sp-proteomics)^30,39^. A custom 300-plex MERSCOPE profiled >1.2 million cells from 6 tissue types, emphasizing cell identity, activation, and receptor-ligand interactions including interleukins, and C-C and C-X-C chemokines, which made up roughly 20% of the panel. After we removed low-count cells, cell types were annotated using TACIT (Supplement Figure 3), and STARComm was then deployed to map receptor–ligand interactions based on spatial proximity and expression specificity in an integrated framework (Supplement Figure 4a)^36^. The highest expressors of these R-L pairs were structural cell types (gland: *ductal epithelial cells*, *acinar cells*; oral mucosae: *keratinocytes*; oral mucosae; both tissue types: *fibroblasts*) (Supplement Figure 4b). We further validated these findings using a single-cell RNA sequencing dataset of the same tissue types, which yielded consistent results, confirming these cellular communication patterns in healthy glands (Supplement Figure 5). In the integrated spatial-transcriptomics dataset, STARComm identified prominent communication pathways primarily involving Acinar cells, which are ubiquitously distributed within the tissue. This suggests a simple and stable interaction network rather than a complex one. Predominant interactions involved signaling pathways such as *TGFA::EGFR*, *HBEGF::EGFR*, *CXCL12::CXCR4*, *CXCL3::CXCR2*, and *EREG::EGFR*, which are known to regulate cellular processes like proliferation, migration, and tissue homeostasis (Supplement Figure 3c). This extensive network of growth factor and chemokine receptor-ligand interactions reflects the dynamic and reciprocal crosstalk that maintains tissue integrity and function in healthy glands, facilitating tissue renewal, immune surveillance, and physiological response to environmental stimuli. These data reveal that even in health, glandular tissues exhibit spatially organized, epithelial-driven signaling networks embedded within the native tissue architecture, thus providing a reference framework to help clinicians detect early immune activation and improve tissue-based diagnostics in conditions like Sjögren’s syndrome, sarcoidosis, and GVHD.

### STARComm Identifies GVHD-Associated Communication MCIMs and Therapeutic Targets

Using high-resolution spatial transcriptomics, a prospective clinical study was conducted to assess whether salivary glands reflect systemic immune dysregulation in chronic GVHD. Minor salivary gland biopsies were collected from 17-chronic GVHD patients (Figure 5a). These minor salivary gland biopsies were processed through our standardized spatial multi-omics pipeline, profiled using Xenium with a targeted 280-plex ISH panel (17 receptor-ligand pairs based on scRNA-seq), and yielded a dataset of 842,154 spatially resolved cells across 36 tissues (8-healthy, 28-GVHD; Figure 5b,c). We then performed an integrated STARComm analysis across all tissues and receptor-ligand pairs to identify shared and divergent spatial communication patterns between healthy and diseased glands. Importantly, 10-GVHD tissues came from patients who later died from GVHD-related complications, enabling STARComm to interrogate spatial signaling changes associated with severe clinical outcomes. This integrated analysis offers a unique opportunity to identify MCIMs with prognostic value and nominate spatially localized, therapeutically targetable pathways relevant not only to salivary gland pathology, but also to systemic GVHD (Figure 5d).

**Figure 5:**
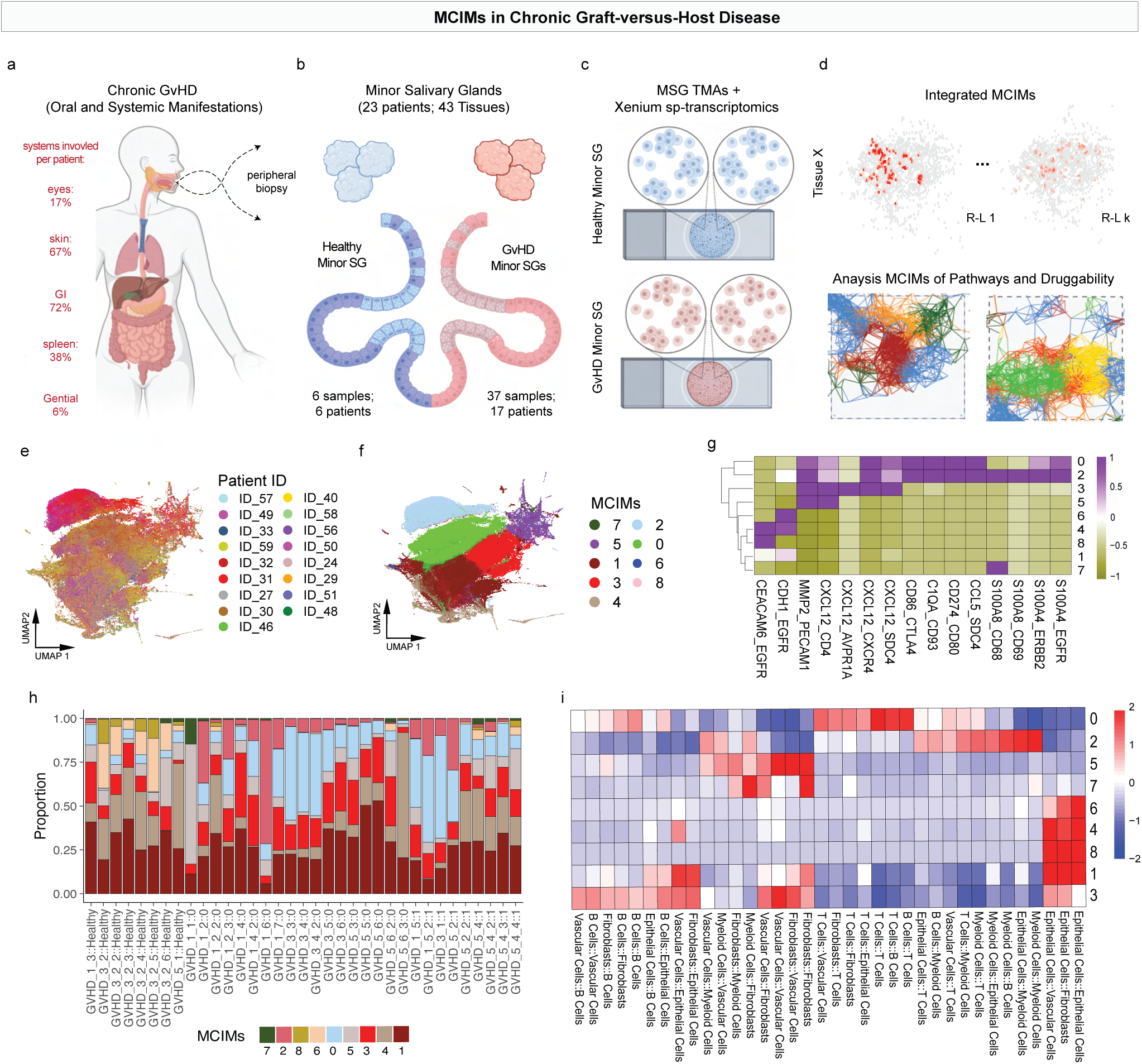
Spatial Analysis of Predicted Receptor–ligand Interactions in GVHD |. **a**, Overview of chronic GVHD manifestations across organ systems in the patient cohort. Percentages indicate the frequency of organ involvement across individuals. **b**, Cohort composition showing 43 minor salivary gland samples collected from 23 patients, including both healthy controls and GVHD cases. **c**, Spatial transcriptomics was performed using the Xenium platform on minor salivary gland tissue microarrays (TMAs), enabling single-cell resolution mapping of gene expression. **d**, Integration of receptor–ligand spatial networks across tissues identified MCIMs based on recurrent cellular interaction patterns (top). These MCIMs were annotated using KEGG and GO enrichment and evaluated for therapeutic relevance using the newly developed spatial Drug2Cell pipeline (bottom). **e–f**, UMAP visualization of bin-level communication densities across all tissues, colored by GVHD status and MCIM membership. **g**, Heatmap showing receptor–ligand density within each MCIM, highlighting signaling pathways associated with disease. **h**, Proportion plot illustrating the distribution of specific MCIMs across 36 tissue samples. **i**, Heatmap depicting enrichment scores of the top interacting cell types within each MCIM.

STARComm detected 9-MCIMs across the 37 tissues (Figure 5e-f; Supplement Figure 6). These MCIMs reflect densely interacting, spatially coherent neighborhoods of immune, stromal, and vascular cells engaged in active signaling. The heatmap of kernel density showing the co-location of multiple cell-cell communication within MCIMs (Figure 5g). The proportion plot of MCIMs illustrating three tissue conditions across 36 tissues, demonstrating the enrichment of MCIM-0 in both GVHD survival and non-survival cases (Figure 5h). We performed TACIT and then visualized data to determine interesting tissue architecture and cell type patterns (Figure 6a). Of particular interest were MCIMs localized to tertiary lymphoid structures (TLS), which were identified by TACIT as regions of high immune cell density. STARComm robustly captured these intralesional communication hubs, most notably MCIM-0, characterized by elevated signaling between B cells, T cells, fibroblasts, and vascular endothelial cells (Figure 5i, Figure 6b-d). Principal component analysis revealed clear separation between healthy and GVHD tissues based on proportion of MCIMs (Figure 6e-f). GVHD tissues showed significantly elevated communication density per unit area (Wilcoxon test, p<0.05), reinforcing the concept of immune overactivation and dysregulated cell–cell crosstalk as central features of GVHD (Figure 6g).

**Figure 6:**
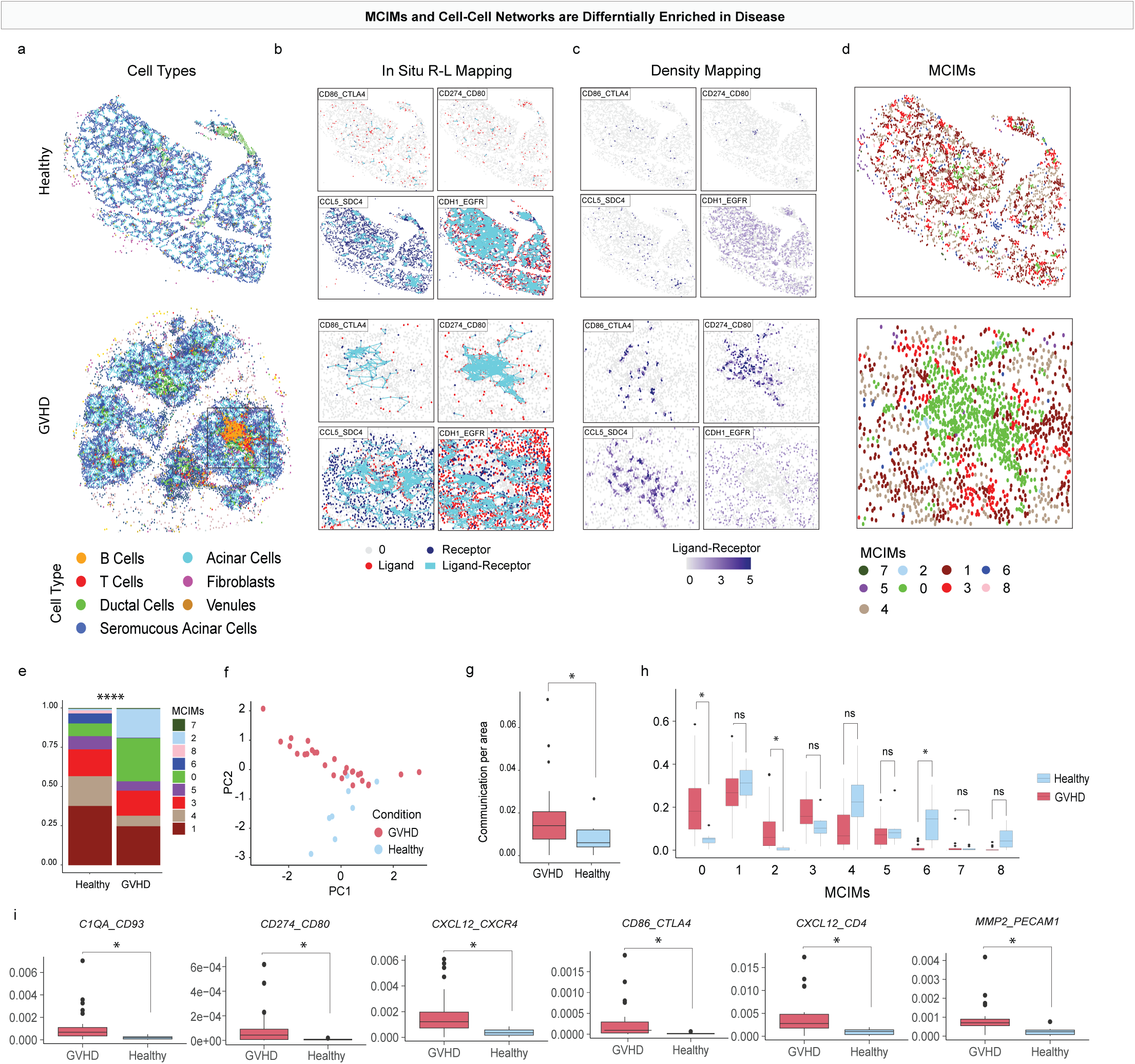
Multicellular Interaction modules (MCIMs) in Chronic Graft-versus-Host Disease (GVHD). | **a**, Visualization of cell type composition in representative healthy and GVHD minor salivary gland samples. Periductal inflammation resembling tertiary lymphoid structures (TLS) is highlighted in the inset. **b**, Spatial maps of four receptor–ligand pairs (CD86–CTLA4, CCL5– SDC4, CD274–CD80, and CDH1–EGFR) using a 50 μm proximity filter. The first three pairs show strong colocalization within TLS regions, while CDH1–EGFR is enriched in healthy tissues but absent from TLS regions. **c**, Receptor–ligand communication density maps highlight spatially enriched zones of signaling activity. **d**, MCIMs identified across individual samples reveal heterogeneous communication profiles within GVHD-affected tissues. **e**, Bar plot showing the distribution of MCIMs in healthy versus GVHD tissues, illustrating disease-associated shifts in multicellular communication (p < 0.01, Wilcoxon test). **f**, Principal component analysis (PCA) of MCIM proportions distinguishes healthy and GVHD tissues based on communication profiles. **g**, Boxplot of overall communication density per tissue area shows significantly higher crosstalk in GVHD tissues (p < 0.05, Wilcoxon test). **h**, Boxplot comparing the proportions of individual MCIMs between healthy and GVHD tissues, identifying significant differences (p < 0.05, Wilcoxon test). **i**, Boxplot comparing receptor–ligand expression levels between GVHD and healthy tissues, highlighting disease-associated upregulation of specific ligand–receptor interactions.

MCIMs-0, -2, and -6 were significantly more frequent in GVHD compared to healthy tissues (Wilcoxon test, p<0.001; Figure 6h), suggesting active roles in disease progression. Key upregulated receptor-ligand pairs within these MCIMs, particularly those associated with tertiary lymphoid structures, included complement activation (*C1QA::CD93*), immune checkpoint regulation (*CD274–CD80*; *CD86::CTLA4*), matrix remodeling (*MMP2::PECAM1*), and immune cell recruitment (*CXCL12::CXCR4*) (Wilcoxon test, p<0.05; Figure 6i). These findings emphasize that MCIMs, not only individual factors, were significantly altered GVHD compared to the healthy This approach gives a chance to more precise in therapeutic application with understanding mechanism through the spatial level in GVHD.

These dysregulated pathways reflect known hallmarks of GVHD pathogenesis and highlight potential therapeutic targets for intervention. Notably, *CD274::CD80* and *CD86::CTLA4* interactions align with emerging checkpoint modulation strategies in autoimmunity and transplantation medicine, suggesting repurposing opportunities^41–43^. The *C1QA::CD93* axis is part of the complement cascade. While direct CD93 targeting is early-stage, complement inhibitors like C1 are under active investigation^44,45^. Furthermore, as *MMP2::PECAM1* is related to fibrovascular activation and architectural reordering, anti-fibrotic agents such as nintedanib may impact related pathways^46,47^. Finally, *CXCL12::CXCR4* governs immune cell retention and tissue homing. CXCR4 inhibitors like *plerixafor* are approved for stem cell mobilization and are being explored for modulating T cell trafficking in GVHD^48–50^. STARComm provides a powerful approach to extract clinically relevant insights from routinely biopsied tissues. By resolving pathogenic cell-cell signaling at the multicellular MCIM level within TLS, it offers a path toward mechanism-based biomarkers and localized therapeutic targeting, with the use of salivary glands

### Spatial Communication MCIMs Predict GVHD Fatality Years in Advance

Given that our GVHD cohort included both survivors and non-survivors, we hypothesized that specific R–L interactions and MCIMs would stratify clinical outcomes and uncover targetable mechanisms of disease progression. Using STARComm, we reanalyzed the 28 GVHD-associated minor salivary gland tissues. Among the MCIMs stratifying clinical outcomes, MCIM-0 emerged as the most differentially enriched between survivors and non-survivors of GVHD (Figure. 7a–c). This MCIMs was characterized by dense B::T cell, T::T cell, and T cell::structural cell interactions, and concentrated 9 of the 17 profiled receptor–ligand pairs—suggesting it represents a dominant communication hub in severe disease. Notably, two key chemokine axes, *CXCL12::CXCR4* and *CCL5::SDC4*, were highly enriched within MCIM-0 and significantly associated with mortality. The strong spatial enrichment of these pathways in fatal disease highlights that only pieces of MCIM-0 serve as a biomarker of immune dysregulation but also focused targets for therapeutic screening. Furthermore, by showing which cell networks were affected in this MCIMs (centrally, T cells were the MCIM-0 drivers; Figure 5i), we gained an opportunity for cell-specific targeting. While these R-L pairs have been previously linked to immune cell recruitment and lymphoid tissue formation, but this is the first demonstration of their spatially resolved, quantitative association with clinical outcomes in GVHD^30,31^.

**Figure 7:**
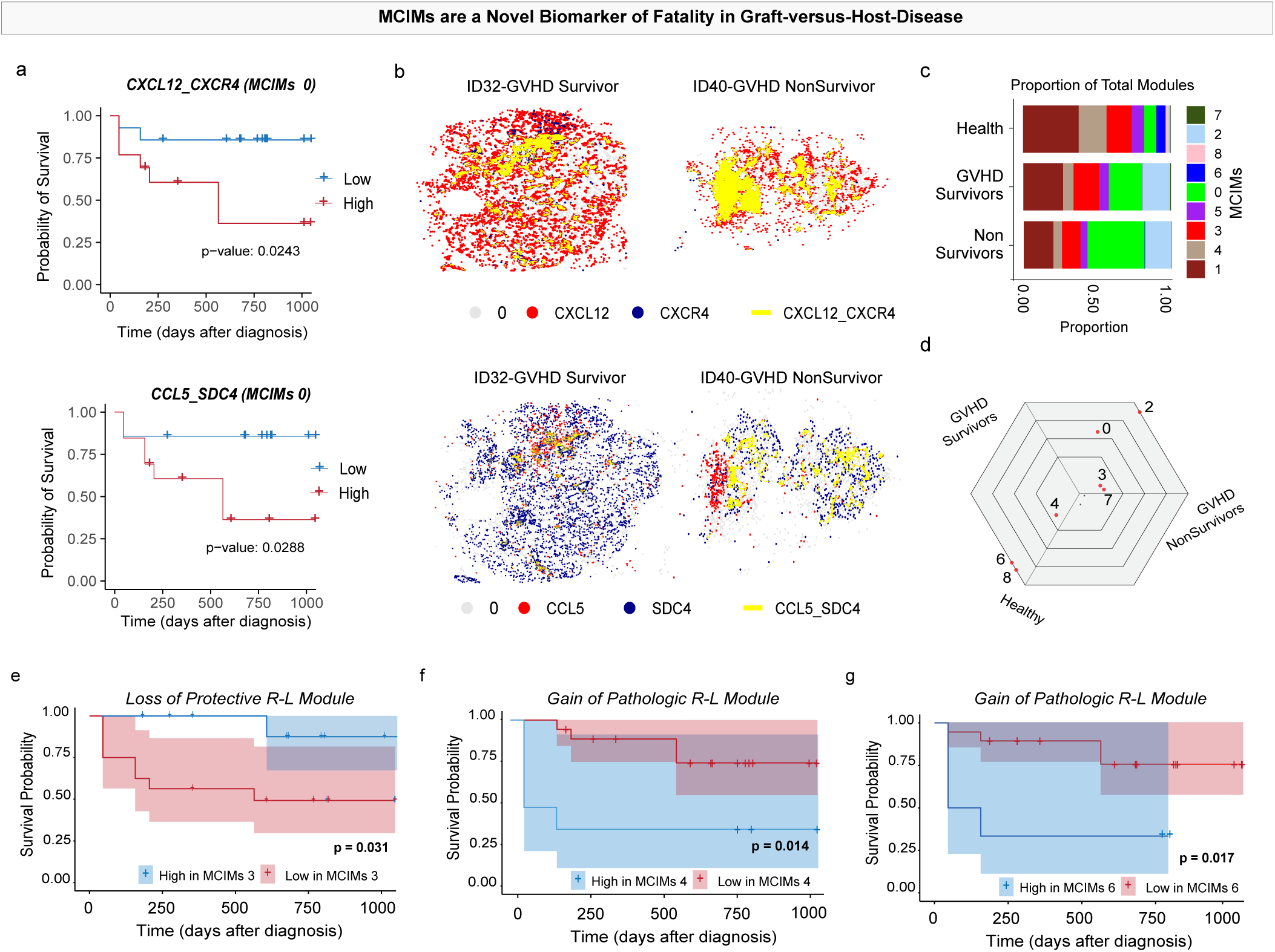
MCIMS Predict Fatality in Chronic GvHD. | **a**, Kaplan–Meier survival analysis shows that higher activity of the receptor–ligand pairs *CXCL12::CXCR4* (p-value = 0.0243, log-rank test) and *CCL5::SDC4* (p-value = 0.0288, log-rank test) within MCIM-0 is significantly associated with reduced survival, indicating their potential contribution to disease progression. **b**, Spatial plots highlight regions of elevated *CXCL12::CXCR4* (top) and *CCL5::SDC4* (bottom) interactions, with these zones corresponding to areas linked to poorer clinical outcomes. **c**, Distribution of MCIM proportions across healthy tissues, GVHD survivors, and non-survivors reveals shifts in communication module dominance across disease states. **d**, Triwise plot comparing MCIM composition among the three groups, with statistically significant differences marked by red dots (ANOVA p < 0.05). Protective MCIMs such as 6 and 8 are enriched in healthy tissues, while MCIMs 0 and 2 are significantly elevated in GVHD patients. **e–g**, Kaplan–Meier curves demonstrate distinct survival associations for specific MCIMs, underscoring their prognostic utility (p-value <0.05, log-rank test).

MCIM comparisons revealed clear communication shifts between healthy controls, GVHD survivors, and non-survivors. MCIMs-6 and -8 predominated in healthy tissues, suggesting homeostatic immune regulation, while MCIMs-0 and -2 enriched across GVHD samples (Figures 7c-d) indicate chronic immune activation through cell networks embedded within tissue structure. Importantly, survival stratification revealed MCIM-3 as pathogenic as patients with lower scores experienced significantly worse long-term outcomes (Figure. 7e). This MCIMs featured receptor-ligand pairs associated with immunoregulation and higher fibroblast-endothelial and B-cell interactions. Conversely, MCIMs-4 and -6 were linked with poorer survival (Figures 7f–g). In a multivariate Cox regression analysis, MCIMs-0, -3, and -6 were the strongest predictors of survival. Elevated MCIM-0 (HR=2.08, *p*=0.038) and MCIM-6 (HR=2.83, *p*=0.003) increased mortality risk, while higher MCIM-3 showed a protective trend (HR=0.51, *p*=0.117). The final model demonstrated strong predictive performance (C-index = 0.80) (Supplement Figure 7), positioning these spatial MCIMs as quantitative biomarkers for GVHD prognosis.

To translate these insights into therapeutic action, we leveraged spatial drug2cell to identify druggable targets within the highest-risk MCIMs^31^. B cells within MCIMs-0 and 6 expressed elevated levels of CD20, nominating Rituximab (anti-CD20 monoclonal antibody) as a high-priority candidate for spatially informed intervention (Supplement Figure 7)^51^. Beyond B cells, these MCIMs also featured signatures of immune checkpoints, fibroblast-endothelial signaling, and chemokine gradients, suggesting that multi-target approaches may be needed. Altogether, spatial MCIMs appear to be mechanistically informative, have the potential to be druggable and/or readouts of druggable activity, and potentially predictive of clinical outcomes as well.

## DISCUSSION

Graft-versus-host disease remains a significant cause of mortality after stem cell transplantation, driven by uncontrolled immune injury to vital organs. While spatial biology has advanced, few studies connect spatially resolved intercellular communication to clinical outcomes. We developed STARComm, a scalable framework for identifying multicellular interaction modules (MCIMs)—spatially co-localized hubs of ligand-receptor signaling reflecting cell-cell communication organization. STARComm integrates cell identity, ligand-receptor expression, and proximity without predefined cell types, prioritizing scalability, efficiency, and prognostic inference.

MCIMs offer a modular, biologically meaningful framework for understanding how multicellular signaling patterns contribute to tissue function and dysfunction. In our analysis, distinct MCIMs mirrored canonical features of GVHD, including persistent immune activation, stromal remodeling, and aberrant lymphocyte trafficking. Notably, MCIM-0 emerged as a high-risk MCIMs consistently enriched in GVHD patients. This MCIMs exhibited dense spatial interactions between B cells and T cells, as well as between T cells and structural or stromal components. It concentrated over half of the high-confidence receptor–ligand pairs profiled in GVHD, including *CXCL12::CXCR4* and *CCL5::SDC4*. These signaling axes are central to immune cell recruitment and positioning, and their therapeutic relevance is underscored by the clinical use of CXCR4 inhibitors like plerixafor. Our findings suggest MCIMs are not just pathological readouts but define actionable communication hubs, informing clinical decisions and targeted interventions. This study first demonstrates spatially defined intercellular communication patterns predict clinical outcomes, opening new avenues for stratified intervention and biomarker discovery.

In the context of GVHD, MCIMs encompassed fibroblasts, lymphocytes, endothelial cells, and epithelial or structural cell types operating in coordinated spatial programs. These findings reinforce the growing appreciation that tissue single-cell pathology emerges more from spatially organized function than from static cell type identity. Unlike conventional histopathological assessment of GVHD, which largely relies on architectural changes and lacks cellular resolution, our approach enables a deeper, cell-level understanding of disease dynamics and immune-stromal interactions in affected tissues^52^.

Analyzing tissues from both healthy individuals and patients with GVHD, we found that MCIMs shift in both abundance and cellular composition during disease progression. MCIM-6 and -8 were largely restricted to healthy tissues, suggesting a role in maintaining homeostatic signaling networks. In contrast, MCIM-0 and -2 were consistently enriched in GVHD, highlighting their involvement in chronic inflammation. MCIM-3 was notable for its protective association: patients with higher MCIM-3 activity demonstrated improved survival, a relationship confirmed by Cox regression modeling. These results build upon earlier spatial biomarker studies—such as those identifying tumor triads or tertiary lymphoid structures—by introducing a multivariate, modular framework that integrates diverse signaling interactions into prognostically meaningful units. Notably, this approach addresses a major gap in the field, as there are currently no reliable predictors of outcomes in patients with chronic GVHD^53^.

To explore therapeutic implications and clinical translation of this workflow, we coupled STARComm with a spatial adaptation of the drug2cell algorithm to identify druggable components within high-risk MCIMs^31^. Within MCIM-0, B cells expressed CD20, suggesting Rituximab as a potential agent to disrupt this fatality-associated MCIMs. Other MCIMs contained immunoregulatory and stromal interactions such as *CD86::CTLA4*, *CD274::CD80*, and *MMP2::PECAM1*. These findings indicate that simultaneous targeting of multiple ligand–receptor pairs may be required to disrupt pathogenic feedback loops. MCIMs thus provide a blueprint for designing multiplexed validation assays, such as spatial proteomics or other smaller plex in situ hybridization assays, that can confirm MCIMs activity in clinical samples and inform therapeutic prioritization, either as standalone tools or as part of emerging combinatorial companion diagnostics in spatial biology.

To validate these findings and correlate oral and other organ MCIMs, we are currently expanding this study to incorporate more subjects with diverse organ involvement to understand within subject. We are also expanding the sp-transcriptomics panel to 1000s of interactions in GVHD and other diseases like polyautoimmunity and pan-airway cancers to support more discovery and subsequent validation. Overall, STARComm facilitates the discovery of spatially organized, and actionable programs that are prognostic. STARComm bridges high-dimensional spatial data to patient outcomes, providing a strong foundation for personalized precision for diagnosis and treatment of a range of human diseases, and GVHD.

## METHODS

### Ethics and Xenium Experiments

This study included all patients diagnosed with oral chronic graft-versus-host disease (cGVHD) between 2017 and 2023 at the Hospital das Clínicas, School of Medicine, University of São Paulo. Patients were included only if histopathological features of cGVHD were present in the oral mucosa or minor salivary glands. Biopsies that had a clinical diagnosis of cGVHD but lacked histopathological confirmation were excluded. The study was approved under IRB protocol 65309722.9.0000.0068 and MTA 5276721.4.0000.0068. The dataset comprises samples from 17 patients, totaling 37 tissue specimens and approximately 842,154 cells. A tissue microarray (TMA) was also constructed, containing 29 cGVHD tissues from patients with follow-up data and 8 healthy minor salivary gland tissues. All samples were derived from FFPE tissue blocks, which were melted and re-embedded into a single recipient block to align with the fiduciary frame of the Xenium slide (10x Genomics). Spatial transcriptomic analysis was performed using the standard 280-plex human breast cancer panel from 10x Genomics, following the manufacturer’s protocol. After processing with the Xenium platform, the slides were sequenced according to the steps described in the Xenium user manual. The slides were then stored in 10% glycerol in PBS for 7 days and subsequently processed using the PhenoCycler Fusion pipeline – Same panel - as first described by our group^36^.

### STARComm Algorithm

#### Input STARComm

##### CELLxFEATURE matrix

Consider the set of genes *G* = {*g*_1_, *g*_2_, … *g*_𝑚_}, which can be divided into ligand genes *L* = {*l*_1_, *l*_2_, … *l*_𝜔_} and receptor genes *R* = {*r*_1_, *r*_2_, … *r*_ẟ_}. The set of genes specific to the spatial transcriptomics panel is then given by *G*_𝑝𝑎𝑛𝑒*l*_ = *G*_*l*_ ∪ *G*_*r*_. The tissue contains a set of n cells, denoted by 𝑆 = {𝑐, 𝑐_2_, … 𝑐_𝑛_}. The gene expression data collected from these cells is represented by a matrix 𝐴 of size 𝑛 𝑥 𝑚, where each element 𝑎_𝑖j_ indicates the number of transcripts detected for gene *j* in cell *i*.

##### Receptor-Ligand list

Receptor-ligand pairs were obtained from public databases such as CellChat and CellTalk, collectively denoted as Π. This set includes pairs Π = {(r_i_, *l*_j_):*r*_𝑖_ ∈ *R*, *l*_j_ ∈ *L*}, where each *r*_𝑖_ is a receptor gene from the set *R*, and each *l*_j_is a ligand gene from the set *L*. The total number of communication pairs is approximately 3,398. The median number of receptors associated with each ligand gene is 3, meaning that, on average, a single ligand interacts with three receptors, while the median number of ligands associated with each receptor gene is 2, indicating that each receptor typically interacts with two ligands.

#### Ligand-Receptor Binding Detection

##### Optimal threshold

A threshold-based approach within the *TACIT* framework is employed to identify cells with positive ligand or receptor expression while reducing background noise. The threshold for each gene *g*_𝑖_ is defined as a function 𝜃(*g*_𝑖_, 𝐴), where 𝐴 is the gene expression matrix.

##### Receptor-Ligand pairs

For each receptor-ligand pair (*l*, *r*) ∈ Π, identify the set of cells expressing positive levels (above the threshold) of the ligand *l* and receptor *r*. Let 𝑆_*l*_ be the set of cells with positive expression of ligand *l*, and 𝑆_*r*_ be the set of cells with positive expression of receptor *r*. Both 𝑆_*l*_and 𝑆_*r*_ are subsets of the overall cells in 𝑆, i.e., 𝑆_*l*_ ⊆ 𝑆 and 𝑆_*r*_ ⊆ 𝑆.

Next, identify all pairs of cells that represent potential receptor-ligand communications. Specifically, for each pair, the ligand-positive cell 𝑐_*l*_ ∈ 𝑆_*l*_ and the receptor-positive cell 𝑐_*r*_ ∈ 𝑆_*r*_ are considered communication pairs if their Euclidean distance satisfies:

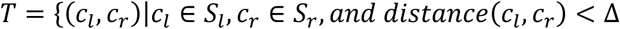

where both 𝑐_*l*_ and 𝑐_*r*_ are cells with positive expression of ligand *l* and receptor *r*, respectively, and their proximity suggests an intercellular communication. It is assumed that each pair (𝑐_*l*_, 𝑐_*r*_) corresponds to a contact point 𝑡 ∈ 𝑇, with a location function 𝑡(𝑐_*l*_, 𝑐_*r*_) defined as the midpoint between the ligand cell 𝑐_*l*_ and the receptor cell 𝑐_*r*_:

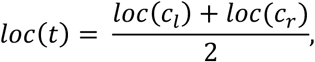

where *l*𝑜𝑐(𝑡) denotes the spatial coordinates of communication 𝑡.

#### Spatial Kernel Density of Ligand Receptors

##### Neighbors of Ligand-Receptor Pairs

The grid is determined by the size of the bins, measured in micrometers (µm). For a grid with bin size 𝐵, the entire tissue is divided into a grid of equally sized intervals, each 𝐵 µm in both the 𝑋 and 𝑌 directions, ensuring each bin is a 𝐵 x 𝐵 µm^2^. Next, for each bin, the function 𝐶𝑜𝑢𝑛𝑡 (𝑐_*l*_, 𝑐_*r*_, 𝐵) calculates the total number of communications for each (*l*, *r*) ∈ Π within the bin 𝐵. This is determined based on the coordinates of the tissue map.

##### Kernel density estimation

Next, Gaussian kernel density estimation (KDE) is employed to estimate the spatial distribution of communication events across the tissue. A higher density value indicates a greater frequency of communication occurring at that location.

*In the 2D case*: Let the location bins be represented by 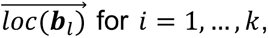, where each point corresponds to a single bin. The receptor-ligand density estimate vector 𝑝(𝒃_𝜏_) at a location 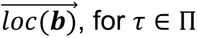, is given by:

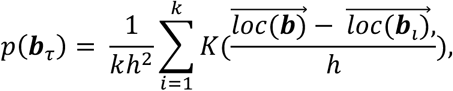

where:

𝑘 is the total number of bins,

ℎ > 0 is the bandwidth parameter,

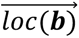 is the location of the bins,

The Gaussian kernel function is: 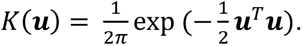.

*In the 3D case*: Similarly, in three dimensions, let the location bins be represented by 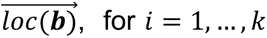, where each point corresponds to a bin in 3D space. The receptor-ligand density estimate vector 𝑝(𝒃_𝜏_) at a location 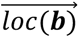 is:

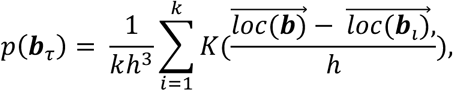

where:

- 𝑘 is the total number of bins,
- ℎ > 0 is the bandwidth parameter,
- 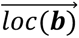 is the location of the bins,
- The Gaussian kernel function is:
- 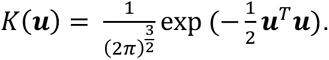

### Identification of Co-localized RL Interactions

Density estimates obtained through Kernel Density Estimation (KDE) are calculated for all receptor-ligand pairs 𝜏, with 𝜏 ∈ Π . These individual density vectors, 𝑝(𝒃_𝜏_), capture the spatial distribution of communication events for each R-L pair across the tissue. By stacking these vectors for all R-L pairs, a matrix 𝐷 is assembled, where each element 𝐷_𝑖j_ represents the density value of the 𝑗-th RL pair (𝜏_j_) at the 𝑖-th bin 𝑏_𝑖_.

The Leiden clustering algorithm is then applied to the matrix 𝐷, grouping bin exhibiting similar patterns of high-density communication into clusters, interpreted as multicellular communication MCIMs (MCIMs). This clustering approach reveals regions with distinct spatial co-location of receptor-ligand activity.

### MCIM Simulation dataset

Let *L* = {*l*_1_, *l*_2_, … *l*_𝜔_} be the set of simulated ligand genes, and *R* = {*r*_1_, *r*_2_, … *r*_ẟ_} be the set of simulated receptor genes. The set of receptor-ligand pairs is defined as

Π = 67r_i_, *l*_j_):*r*_𝑖_ ∈ *R*, *l*_j_ ∈ *L*},

Each *r*_𝑖_ ∈ *R* is a receptor gene, and each *l*_j_ ∈ *L* is a ligand gene. The set of cells in the simulated tissue sample is denoted by 𝑆 = {𝑐_1_, 𝑐_2_, … 𝑐_𝑛_}, with |𝑆| = 𝑛. Each cell 𝑐_𝑖_ ∈ S is assigned to a ground truth multicellular communication MCIMs (MCIM), which serves as the reference label for clustering evaluation.

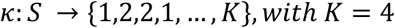

Where 𝜅(𝑐_𝑖_) = 𝑘 indicates that cell 𝑐_𝑖_ belongs to the 𝑘^𝑡ℎ^ MCIM. For each MCIM, a subset of receptor-ligand pairs 67*l*_𝑖_, *r*_j_)h ⊂ Π is randomly selected to be enriched within that MCIM. Next, the pairwise Euclidean distance between cells ci and c j is calculated as:

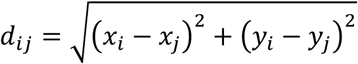

where (𝑥_𝑖_, 𝑦_𝑖_) are the spatial coordinates of cell 𝑐_𝑖_. Edges (𝑐_𝑖_, 𝑐_j_) representing potential communication links are sampled with a probability that decreases with the spatial distance between cells. Specifically, for each pair (𝑐_𝑖_, 𝑐_j_), the probability of an edge is modeled as:

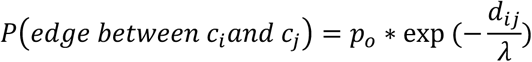

where 𝑝_𝑜_ is a base probability, and 𝜆 is a decay parameter that controls how rapidly the probability decreases with distance. In this model, pairs separated by less than 50µm have a higher probability of forming an edge, while pairs beyond 50µm retain a non-zero probability that diminishes exponentially with increasing distance. For each sampled edge (𝑐_𝑖_, 𝑐_j_) that carries communication, ligand or receptor expression levels are assigned. If cell ci carries a ligand/receptor, its expression level for that ligand/receptor gene 𝑖 is sampled from a Negative Binomial distribution with a high mean:

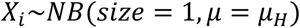

If the cell does not carry the ligand or receptor, the expression is sampled with a baseline mean:

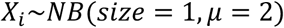

Hence, in this simulation, the density of cell-cell communication can be regulated by modifying the base probability 𝑝_𝑜_, the number of receptor-ligand pairs (by adjusting ω or 𝛿), and the high-expression mean 𝜇_𝐻_.

### Spatial Autocorrelation Analysis using Moran’s I

To evaluate whether MCIMs are spatial co-location in the tissue, Moran’s I statistic is computed for each MCIM across the spatial domain. Let 𝑆 = {𝑐_1_, 𝑐_2_, … 𝑐_𝑛_} denote the set of cells, where each cell 𝑐_𝑖_ has spatial coordinates (𝑥_𝑖_, 𝑦_j_). For a given MCIM 𝑘, define a binary indicator variable:

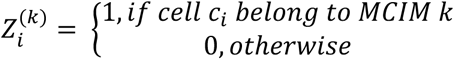

Let 𝑊 = [𝑤_𝑖j_] be a spatial weights matrix, where 𝑤_𝑖j_ reflects the spatial proximity or connection strength between cells 𝑐_𝑖_ and 𝑐_j_, normalized so that ∑_𝑖,j_ 𝑤_𝑖j_ = 1. The Moran’s I statistic for MCIM 𝑘 is calculated as:

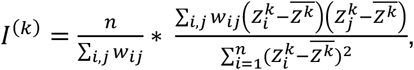

Where:

- 𝑛 is the total number of cells,
- 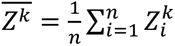 is the mean of 𝑍^𝑘^

Hypotheses:

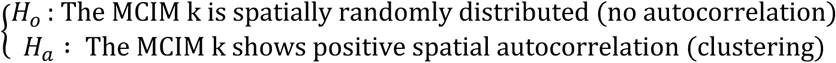

Significance of the Moran’s I value is evaluated through permutation tests, where the labels 𝑍^𝑘^are randomly shuffled many times to generate a null distribution. The resulting p-value indicates whether the clustering of MCIM 𝑘 is statistically significant.

### Calculation of the area of the tissues

To calculate the tissue area based on coordinate points in R, you can use the chull() function to identify the convex hull of the points. First, pass your x and y coordinates to chull() to obtain the indices of the points forming the convex hull. Next, extract the corresponding hull coordinates and use a custom function implementing the shoelace formula—such as the polyarea() function provided—to compute the polygon’s area. This approach gives you the approximate tissue area enclosed by the outermost points.

### Benchmarking with other methods

We compared our proposed method with two existing spatial cell-cell communication methods and graph-based clustering methods, namely COMMOT, SpatialDM, and Leiden. The code for COMMOT, SpatialDM, and Louvain methods are publicly available for reproducibility and comparison purposes.

### COMMOT

COMMOT is a spatial cell-cell communication algorithm that uses collective optimal transport theory to infer ligand-receptor interactions. This approach preserves the comparability of distributions, ensures the total signaling does not exceed the available ligand or receptor amounts, enforces spatial limits on signaling, and accommodates multiple competing species. By jointly optimizing the total transported mass and ligand-receptor coupling, COMMOT provides a comprehensive inference of cell-cell communication compared to existing optimal transport methods. To run COMMOT, we follow the steps from the official GitHub repository (https://github.com/zcang/COMMOT): set the distance threshold to 50 µm, and use the output from the analysis for further clustering. We take the commot-user_database-sum-receiver and commot-user_database-sum-sender files as cell by feature matrix and perform Leiden clustering on the combined matrix and analyze the resulting clusters against the ground truth.

### SpatialDM

SpatialDM (Spatial Direct Messaging, or Spatial co-expressed ligand and receptor Detected by Moran’s bivari- ant extension) is used to identify spatial co-expression between ligand and receptor pairs through a bivariate Moran’s statistic, which assesses spatial association. The workflow begins by assigning spatial coordinates to adata2.obsm[’spatial’] from the dataset. A weight matrix is constructed using sdm.weight_matrix(), with parameters l = 1.2 and a cutoff of 0.2, via an RBF kernel to model spatial relationships. Gene interaction data, including ligand-receptor pairs, are stored in adata2.uns under entries like ’geneInter’, ’receptor’, and ’ligand’. Global spatial association is then evaluated with sdm.spatialdm_global(), testing overall Moran’s association across all spots. Significant ligand-receptor pairs are identified using sdm.sig_pairs() with permutation testing and FDR correction. Local co-localization is assessed with sdm.spatialdm_local(), followed by filtering significant spots via sdm.sig_spots(). These significant spots are binarized based on a threshold, and spatial patterns are analyzed with SpatialDE through SpatialDE.run(), executed within a limited BLAS thread context using threadpool_limits(). The resulting spatial patterns are clustered using kmeans(), with a specified number of clusters (e.g., 4), and these labels are used for downstream spatial clustering, such as Leiden clustering, to identify spatially co-localized cell populations.

### ExpressionOnly

Leiden clustering is a popular unsupervised method for identifying clusters in scRNA datasets. Originally developed for community detection in networks, it optimizes modularity to partition data into clusters, effectively distinguishing cell populations based on gene expression profiles. To perform Louvain clustering, we first normalized the data using z-score normalization, then scaled it to give equal weight to all features. We applied UMAP for dimensionality reduction on the top 10 principal components, allowing visualization in a lower-dimensional space. Clustering was then carried out on the UMAP coordinates with a resolution that can gave 4 clusters.

### Performance metrics

The clustering performance was evaluated using the Adjusted Rand Index (ARI), which quantifies the similarity between the predicted clusters and the simulated true labels in the simulated data. The ARI was calculated using the aricode package version 1.0.3. An ARI score of 1 indicates perfect agreement, while scores closer to 0 or negative values suggest poor or no agreement. This metric was used to compare the effectiveness of different clustering methods against the simulated ground truth.

### Statistical Analyses

Statistical analyses and figure generation were performed using R (version 4.3.0). When comparing two groups, a Student’s t-test was employed if data met the assumption of normality; otherwise, the non-parametric Wilcoxon rank-sum test was utilized. For comparisons involving more than two groups, analysis of variance (ANOVA) was conducted, followed by post-hoc tests if significant differences were observed. To correct for multiple testing, the false discovery rate was controlled using the Benjamini-Hochberg procedure. Statistical significance was defined as P < 0.05, or an adjusted P < 0.05 for multiple comparisons.

## CODE AVAILABILITY

The code used to develop the STARComm, perform the analyses and generate results in this study is publicly available and has been deposited in https://github.com/huynhkl953/STARComm.

## Contributions

For this study, KMB and JL conceptualized the project. JL, and KLAH designed the algorithm. KLAH implemented the algorithm and conducted benchmarking experiments. BFM performed manual assessment of GVHD clinical outcome. DDS, GF, LAVSJ, LFFDS, and VGR, supported the recruitment of patients and collected data. KLAH, XZ, BFM, DDS, GF, LAVSJ, LFFDS, VGR, KMB, and JL performed experimental and/or bioinformatic analysis that supported project development. JL, KMB, KLAH, and BFM wrote the original draft; KLAH, XZ, BFM, DDS, GF, LAVSJ, LFFDS, VGR, KMB, and JL critically reviewed and edited the final manuscript.

## Acknowledgements

ADA Science & Research Institute (startup funds & Volpe Research Scholar Award, K.M.B.); and the Department of Oral and Molecular Craniofacial Biology, Philips Institute for Oral Health Research (startup funds, K.M.B.). The work also benefited from the VCU Wright Regional Center for Clinical & Translational Science (CCTS) Clinical and Translational Science Award (CTSA) UM1TR004360 to JL and NIH-NCI Cancer Center Support Grant P30 CA016059 to JL.

---

**Supplement Figure 1:**
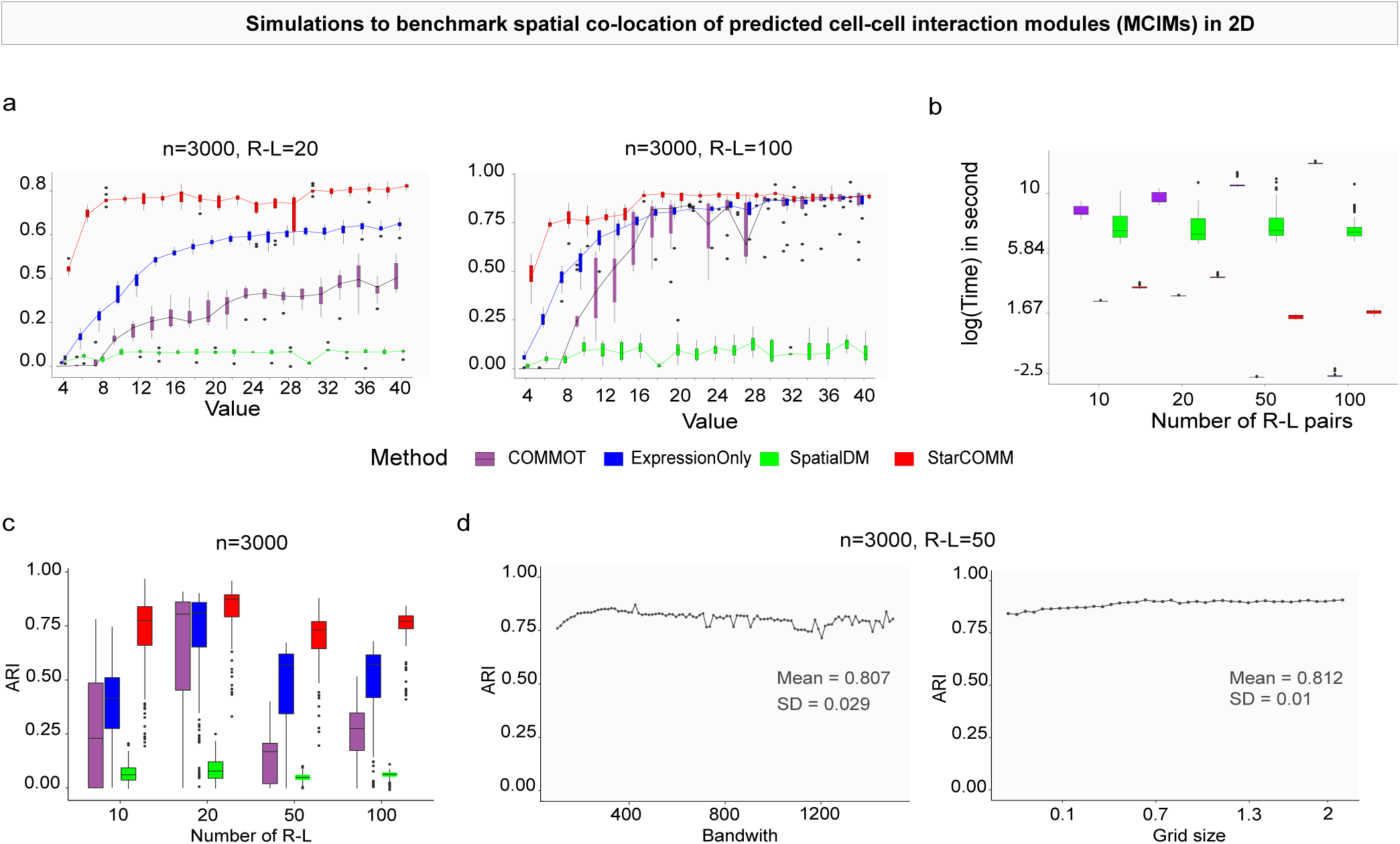
**a**, Simulated datasets were generated to model spatially localized cell– cell communication events, with the number of receptor-ligand pairs set at 20 and 100. **b**, The runtime of each method, demonstrating that both STARComm and ExpressionOnly have the shortest execution times compared to COMMOT and SpatialDM. **c**, The ARI scores across all methods, indicating that STARComm’s performance is unaffected by the number of receptor-ligand pairs. **d**, Compares STARComm results using different parameters, such as small bandwidth and grid size.

**Supplement Figure 2:**
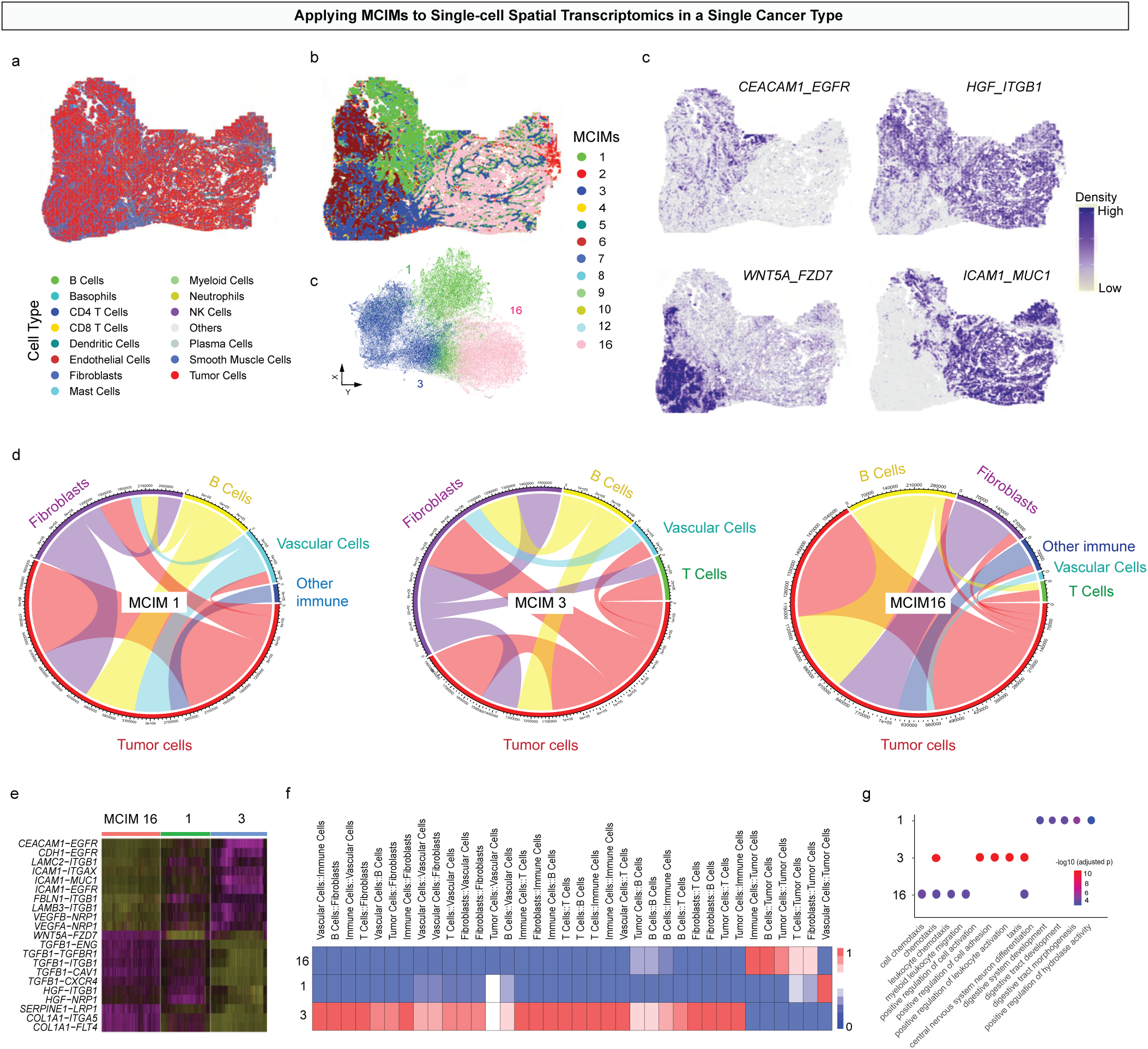
**a**, shows the spatial map highlighting tumor cells in red within the tissue. **b**, depicts the three distinct MCIM regions, each characterized by unique patterns of cell–cell co-location and communication. **c**, displays the density maps of communication enrichment within each MCIM, indicating regions of active signaling. **d**, illustrates the receptor–ligand interactions between MCIMs 1, 3, and 16, highlighting the pathways involved in tumor growth, immune regulation, and cell migration. **e**, lists the top enriched receptor–ligand pairs specific to each MCIM, with pathways such as CEACAM1–EGFR, ICAM1–MUC1, and WNT5A–FZD7. **f**, visualizes cell type communication networks within and between regions, revealing how different cellular populations interact across the tumor microenvironment. Finally, **g**, presents KEGG pathway enrichment analysis for MCIMs 1, 3, and 16, highlighting active biological processes such as tissue development, immune response, and cell migration that underscore the functional roles of these regions in tumor progression.

**Supplement Figure 3:**
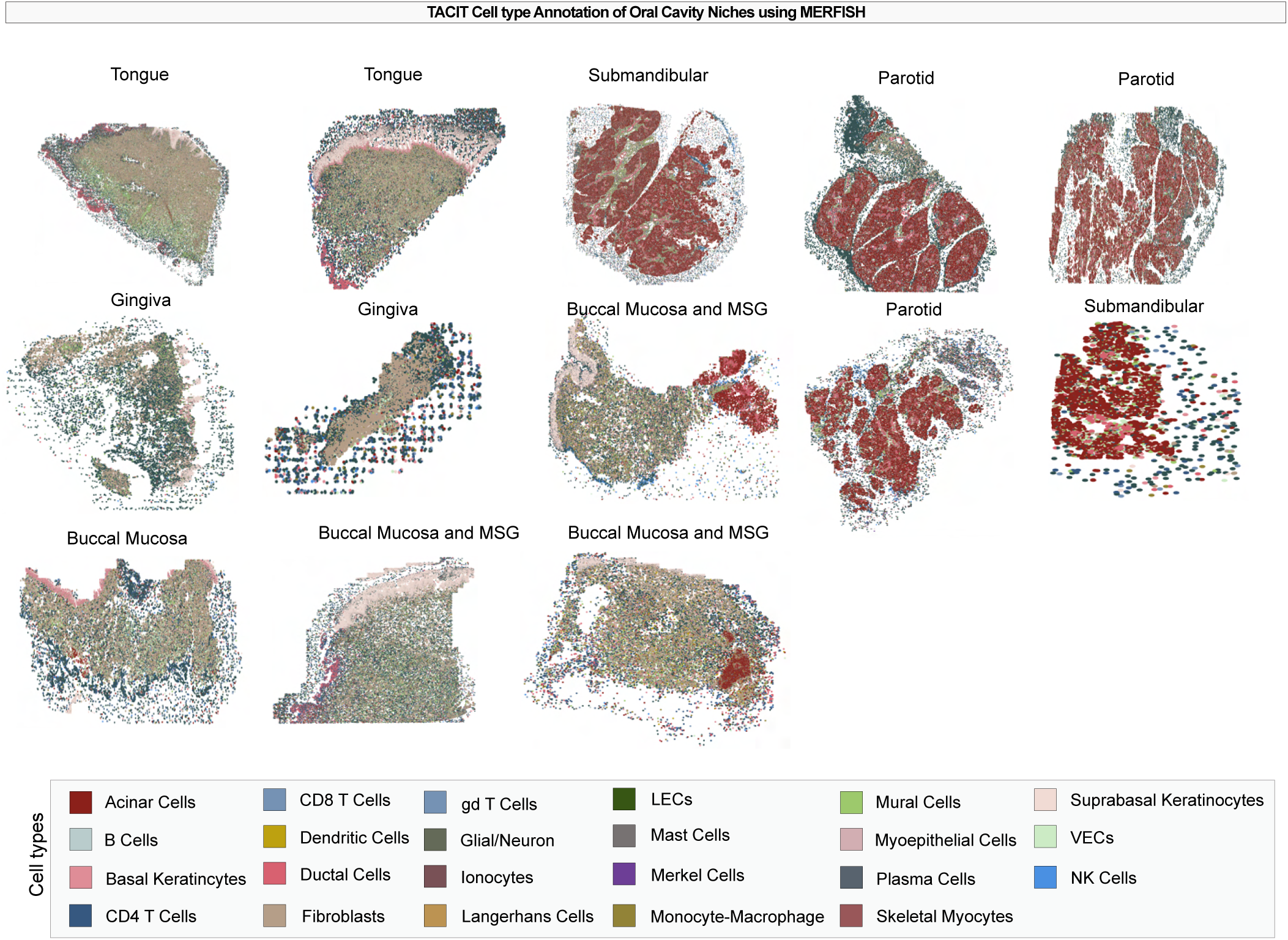
TACIT Cell type Annotation of Oral Cavity Niches using MERFISH.

**Supplement Figure 4:**
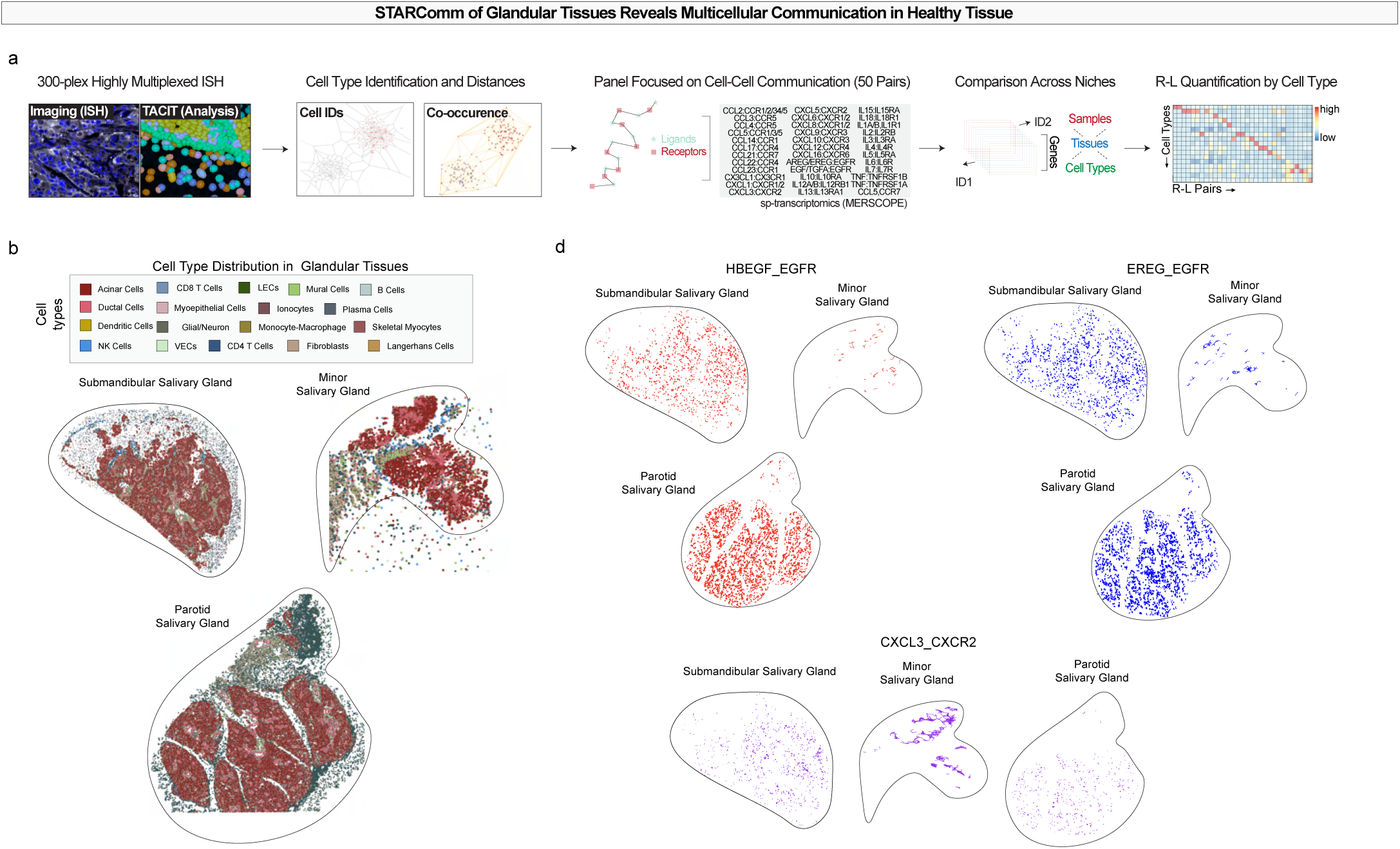
**a**, Multiplexed in situ hybridization imaging was performed using a 300-gene MERSCOPE panel. Cell identities were assigned using TACIT based on transcript expression. Receptor and ligand coordinates were mapped within annotated cell types using the CellChat receptor–ligand database. STARComm was then used to compare receptor–ligand interaction patterns between mucosal and glandular oral niches, followed by clustering of interaction modules across tissue types. **b**, Voronoi plot showing MERSCOPE-derived cell type annotations, with diverse epithelial, stromal, immune, and vascular populations serving as spatial anchors for receptor–ligand interaction analyses. **c**, Heatmap of cell type communication within each MCIMs in glands. **d**, In glandular tissues (parotid, minor salivary, and submandibular glands), 10 MCIM clusters were identified. MCIMs 2 and 13 exhibited unique ligand–receptor interaction profiles not observed in other modules. Spatial network analysis revealed that MCIMs 1 and 4 served as central hubs with high levels of interconnectivity. **e**, Dotplot of communication within each MCIMs in glands.

**Supplement Figure 5:**
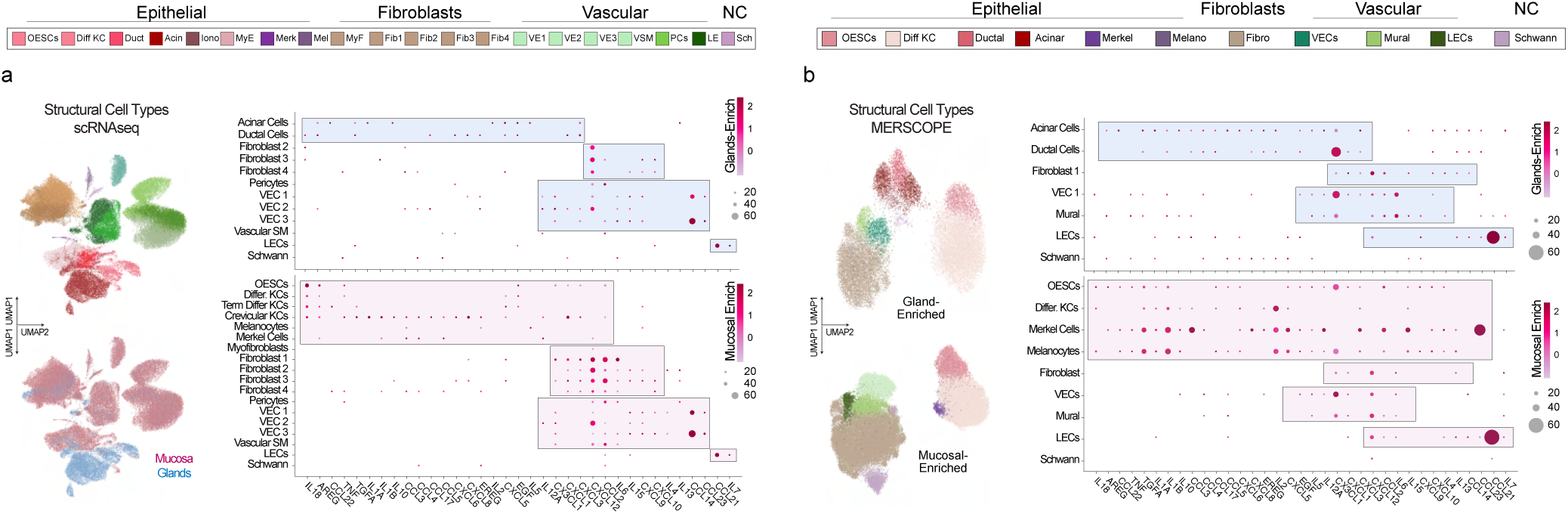
scRNAseq analysis revealed structural ligand signatures enriched exclusively in one structural compartment—either epithelial or stromal—within glands and mucosa and in distinct cell types.

**Supplement Figure 6:**
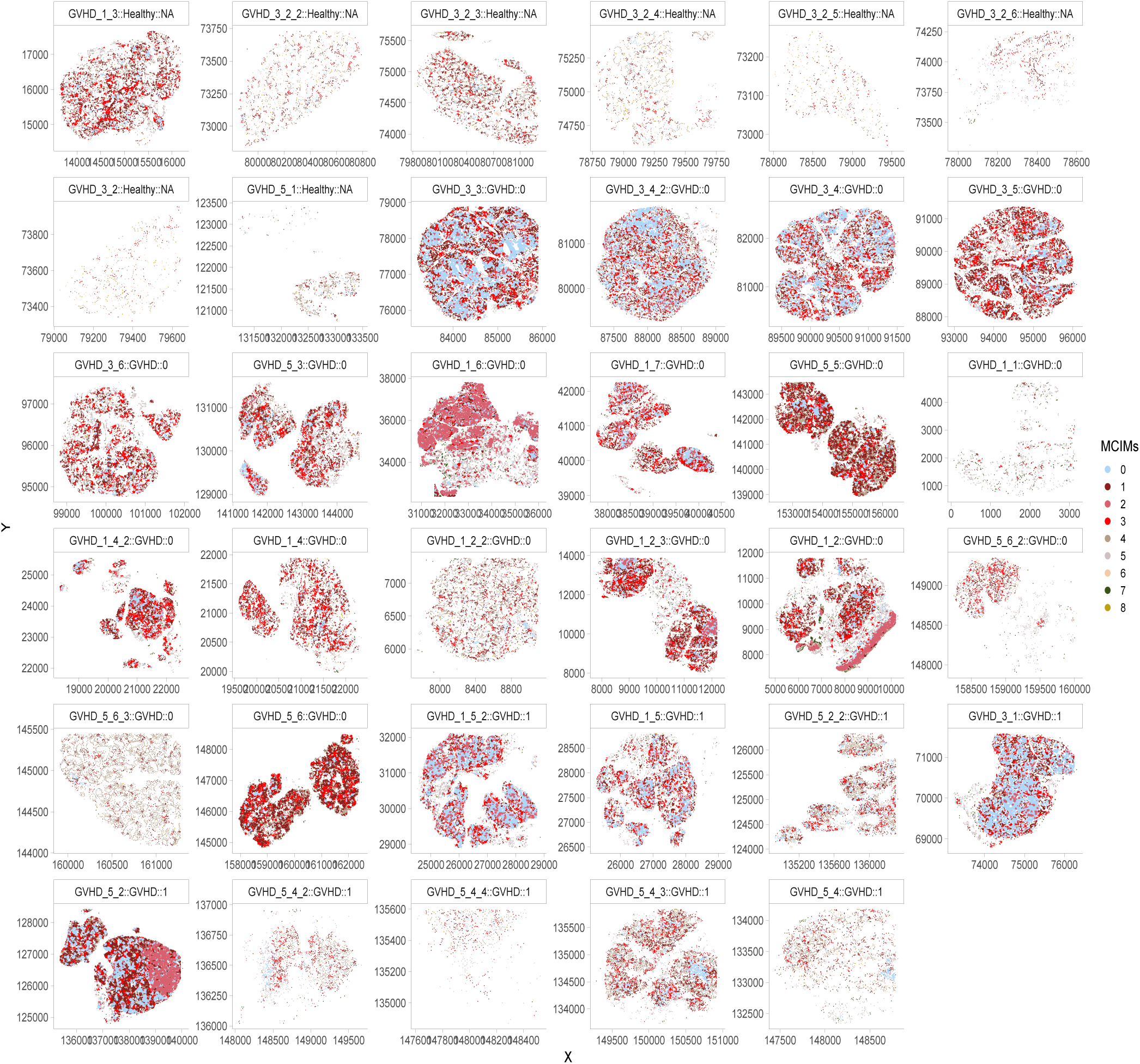
Spatial MCIMs across 36 tissues in GVHD cohorts with survival outcome.

**Supplement Figure 7:**
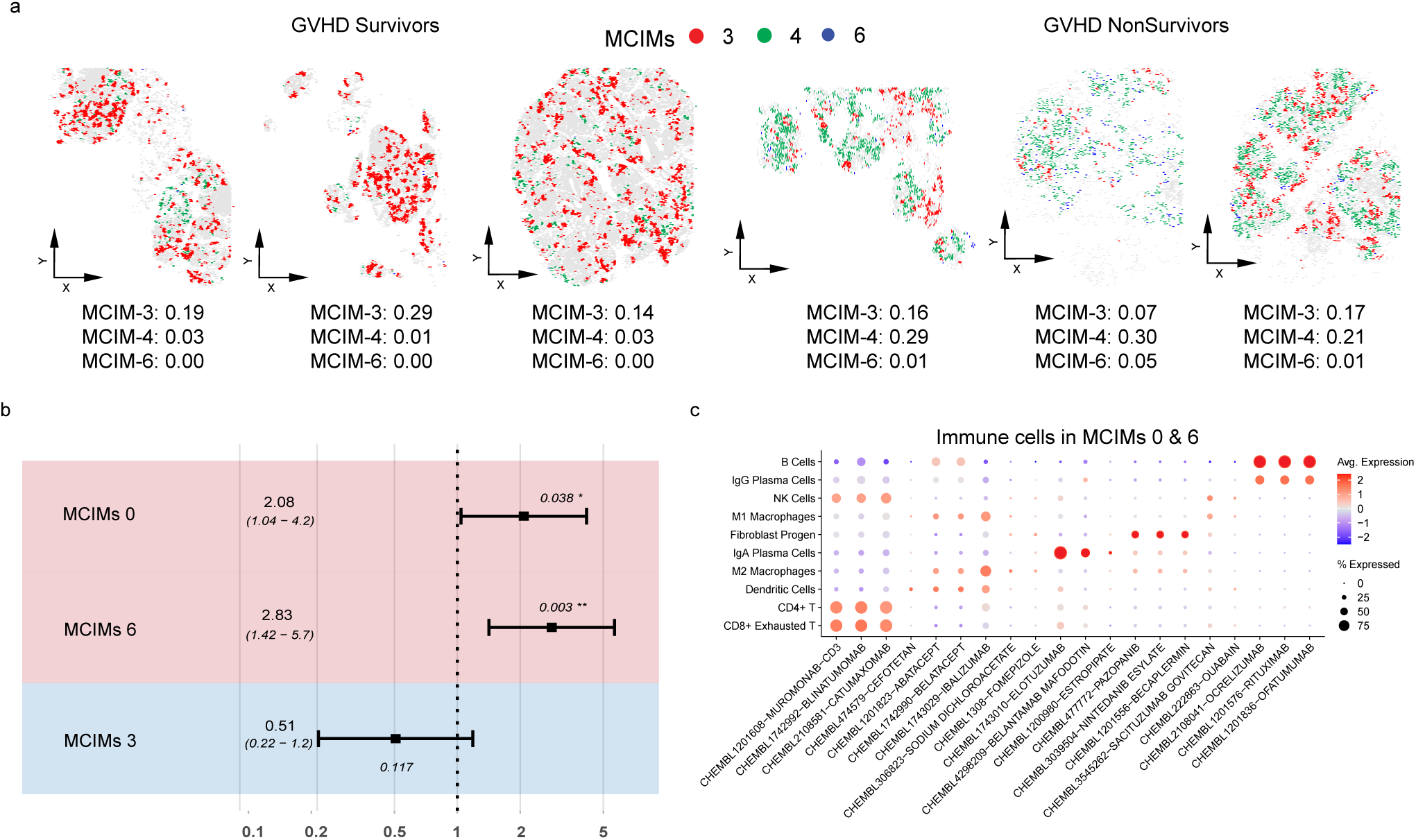
**a**, Examples of spatial MCIM-3, -4, and -6 in GVHD survivors and non-survivors. **b**, The optimal Cox regression model identified through backward selection, highlighting MCIM-0, -6, and -3. **c**, The Spatial drug2cell analysis focusing on immune cell types within MCIM-0 and -6.

